# Engineering affinity-matured variants of an anti-polysialic acid monoclonal antibody with superior cytotoxicity-mediating potency

**DOI:** 10.1101/2025.02.12.637914

**Authors:** Weiyao Wang, Mehman Bunyatov, Natalia Lopez-Barbosa, Matthew P. DeLisa

## Abstract

Monoclonal antibodies (mAbs) that specifically recognize cell surface glycans associated with cancer and infectious disease hold tremendous value for both basic research and clinical applications. However, high-quality anti-glycan mAbs, especially those with sufficiently high affinity and specificity, remain scarce, highlighting the need for protein engineering approaches based on rational design or directed evolution that enable optimization of antigen-binding properties. To this end, we sought to enhance the affinity of a polysialic acid (polySia)-specific antibody called mAb735, which was raised by animal immunization and possesses only modest affinity, using a combination of rational design and directed evolution. The application of these approaches led to the discovery of affinity-matured IgG variants with up to ∼7-fold stronger affinity for polySia relative to the parental antibody. The higher affinity IgG variants were observed to opsonize polySia- positive cancer cells more avidly, which in turn resulted in significantly greater cytotoxicity as determined by both antibody-dependent cell-mediated cytotoxicity (ADCC) and complement-dependent cytotoxicity (CDC) assays. Collectively, these results demonstrate the effective application of both rational and random molecular evolution techniques to an important anti-glycan antibody, providing insights into its carbohydrate recognition while at the same time uncovering variants with greater therapeutic promise due to their enhanced affinity and potency.

## INTRODUCTION

Monoclonal antibodies (mAbs) are the fastest-growing class of biological therapeutics and have revolutionized the treatment of various hematologic and solid malignancies ^1–4^ as well as infectious diseases ^5–8^. While most clinically approved mAbs are directed against proteins, in recent years carbohydrate chains known as glycans have gained increasing attention as therapeutic targets ^9–11^. The promise of glycans as anti-cancer targets derives from the observation that cell-surface glycosylation patterns change during malignant transformation, leading to abnormal tumor-associated carbohydrate antigens (TACAs) that are abundantly and selectively expressed on cancer cells ^12–15^. In the context of infectious disease, the glycans present on the surfaces of bacterial, viral, and fungal pathogens are attractive targets because they are often distinct from those produced by healthy human cells ^16, 17^. Accordingly, mAbs that specifically recognize these cancer- and infectious disease-associated glycans hold enormous clinical value. For example, dinutuximab (Unituxin) and naxitamab (Danyelza), both of which target the ganglioside GD2, have been approved by the FDA for treatment of high-risk pediatric neuroblastoma and are the first anti-TACA mAbs to be successfully translated to the clinic.

Despite the uptick in the number of anti-carbohydrate mAbs undergoing clinical evaluation ^10^, their binding properties are often suboptimal compared to antibodies targeting proteins. In general, anti-glycan mAbs exhibit affinities that are 1,000 to 100,000 times lower than the affinities of anti-protein or anti-peptide antibodies for their antigens ^9, 11^ and suffer from widespread specificity problems as judged from the high number of existing anti-glycan mAbs that cross-react with other glycans ^18^. There are several reasons for the relatively low affinity and high non-specific binding of anti-glycan mAbs derived from an immunized host. For one, unlike protein antigens, most carbohydrates are T cell-independent antigens, which trigger B-cell responses that lack affinity maturation and are biased toward the production of IgM ^19–21^. Furthermore, anti- carbohydrate immune responses generate antibodies from a limited repertoire of variable (V) region genes with restricted gene pairing ^22–26^. Collectively, these phenomena lead to the expression of essentially germline antibody sequences characterized by low affinity and broad specificity ^11, 27^.

To overcome these binding liabilities, it is necessary to generate mutants of pre- existing anti-glycan antibody scaffolds with enhanced affinity, selectivity, and specificity. A variety of protein engineering approaches based on rational design or directed evolution have proven useful for optimizing the antigen-binding properties of antibodies. A common workflow involves screening combinatorial libraries of recombinant antibody genes– typically in the single-chain fragment variable (scFv) or fragment antigen-binding (Fab) format–using display technologies such as yeast surface display and filamentous phage display ^28, 29^. However, while these strategies have met widespread success in the context of anti-protein and anti-peptide antibodies, their implementation for anti-glycan antibodies has significantly lagged and yielded mixed outcomes ^27, 30–37^. For example, Brummel et al. constructed 90 mutants of a Fab antibody specific for *Salmonella* serogroup B O- polysaccharide (O-PS) by site-directed mutagenesis of the heavy chain complementarity determining region 3 (CDR H3); however, none of the tested mutants showed improved binding affinity for the O-PS antigen ^37^. Even in cases where binding affinity was improved, maintenance of antigen specificity has proven challenging, as exemplified by the phage display-based isolation of an affinity-matured scFv antibody against GD2, which exhibited 19-fold higher affinity for the target ganglioside but also evolved strong cross-reactivity to other related ganglioside structures that was not observed with the parental scFv antibody ^36^. Several other studies also reported that affinity maturation of anti-glycan antibodies was accompanied by altered specificity ^31, 34^.

Collectively, these issues provide a rationale for the wider application of protein engineering tools to pre-existing anti-glycan antibodies. To this end, we focused on an existing anti-glycan IgG2a antibody named mAb735 that was developed in an autoimmune mouse strain and specifically recognizes a homopolymer of α2,8-linked *N*- acetylneuraminic acid (Neu5Ac) sialic acid residues called polysialic acid (polySia) ^38^. PolySia occurs as a terminating structure on *N*-linked glycans associated with the neural cell adhesion molecule (NCAM) in vertebrates and as a capsular polysaccharide structure on the surface of bacterial pathogens that cause meningitis and sepsis ^39^. In vertebrates, polySia is an oncofetal antigen that has significantly reduced expression in healthy adults but is aberrantly re-expressed during progression of several malignant human tumors, including small-cell lung cancer (SCLC), non–small cell lung cancer (NSCLC), neuroblastoma, and pancreatic cancer, among others ^40–43^. Notably, among high priority cancer antigens, polySia was the second ranked TACA (after GD2) in a National Cancer Institute pilot project ^44^.

Here, we performed a combination of rational design and directed evolution of an anti-polySia scFv derived from mAb735 (scFv735). Specifically, we used structure-guided site-directed mutagenesis (SDM) to exhaustively probe the binding contributions of all CDR residues that are observed to interact with polySia in the solved crystal structure of the antibody-glycan complex ^45^. In parallel, we used yeast surface display to screen combinatorial libraries of scFv735 variants in which random mutations were introduced either throughout the entire gene or within the CDRs only. These protein engineering approaches enabled identification of residues both within and outside the paratope that are essential for polySia recognition and that increased affinity for polySia by up to ∼4- and ∼7-fold in the scFv and IgG formats, respectively. The higher affinity IgG variants were found to bind polySia-positive tumor cells more avidly and exhibited significantly greater tumor cell killing as determined by both antibody-dependent cell-mediated cytotoxicity (ADCC) and complement-dependent cytotoxicity (CDC) assays. Taken together, these results demonstrate the successful application of protein engineering approaches to an important anti-glycan antibody, resulting in detailed molecular insights into carbohydrate recognition and discovery of several variants that hold therapeutic potential due to their enhanced affinity and potency.

## RESULTS

### Structure-guided identification of residues that contact polySia antigen

Akin to other anti-glycan mAbs derived from animal immunization, mAb735 is not substantially different from germline gene segments and exhibits a modest affinity (*K*_D_ ≈ 4 μM for scFv and Fab derivatives) for long-chain polySia having a degree of polymerization (DP) >40 ^43, 46^. The determination of contact residues from high-resolution antibody-antigen complex structures provides information that can be used for paratope mapping as well as guiding the affinity maturation process ^47^. Accordingly, we took advantage of the published crystal structure of scFv735 in complex with octasialic acid ^45^, which revealed extensive interactions across all CDRs in the variable heavy (V_H_) and variable light (V_L_) domains except for CDR L3 (**Fig. 1a**). A total of six residues were observed to directly interact with the octasialic acid ligand. These direct interactions were formed between the hydroxyl groups of the sialic acids and either the hydroxyl groups in the side chains of Tyr-37 (in CDR L1), Tyr-159 and Tyr-160 (both in CDR H1), and Tyr-179 (in CDR H2) or the polar side chains of Arg-55 (in CDR L2) and Asp-232 (in CDR H3). In addition to direct contacts, we also observed a water-mediated hydrogen bond network that involved all CDRs except for CDR L3 and stabilized octasialic acid binding (**Fig. 1b**). This network was composed of ten residues that indirectly interacted with octasialic acid including Arg- 55 and Asp-232, which also directly contacted the bound ligand. Altogether, 14 residues comprising the paratope were identified by this analysis, with the majority occurring within CDRs of the V_H_ domain.

**Figure 1.**
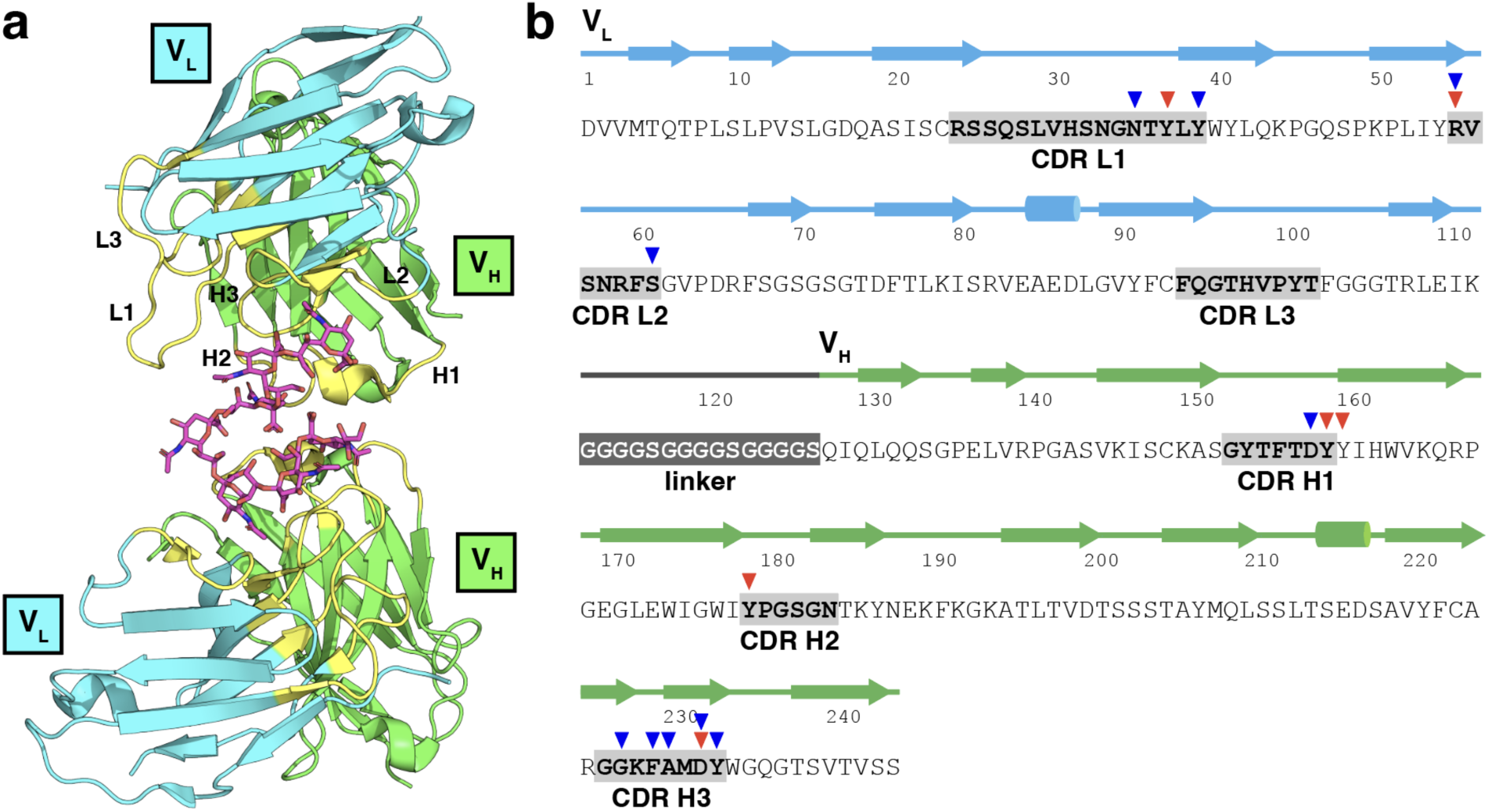
Structural basis for polySia binding by scFv735. (a) Crystal structure of an anti-polySia scFv derived from mAb735 (scFv735) in complex with octasialic acid as reported by Nagae et al.^45^ Protein and carbohydrate are shown as ribbon and rod models, respectively. The variable heavy (VH) domain is colored in green, variable light (VL) is colored in cyan, CDRs H1-H3 and L1-L3 are colored in yellow, and carbohydrate is colored in purple. (b) Amino acid sequence of scFv735 depicted in (a). The secondary structural elements of the VL (cyan) and VH (green) domains, as well as the 15-residue (Gly4Ser)3 flexible linker (dark grey) that connects the VL and VH domains are indicated above the sequence, with α-helices and β-strands denoted by cylinders and arrows, respectively. The CDRs (bold font and gray shaded boxes) were determined using an immunoinformatic analysis tool called antibody region-specific alignment (AbRSA) (http://cao.labshare.cn/AbRSA).^48^ Red and blue triangles above the sequence denote the residues that make direct and water-mediated indirect interactions, respectively, with the polySia antigen based on the solved crystal structure.^45^

### Comprehensive mutational scanning of direct and indirect contact sites

In the context of protein antigens, antibody paratopes often exhibit a considerable degree of plasticity in that multiple amino acid substitutions can be tolerated in these regions and occasionally improve affinity ^49–51^. Therefore, to enable a comprehensive analysis of the plasticity of the paratopic amino acids and potentially uncover substitutions that enhance affinity, we subjected all direct and indirect contact residues in scFv735 to site-saturation mutagenesis (SSM). SSM involves replacing a codon of a gene with codons for all the other 19 amino acids and is commonly implemented using PCR amplification with degenerate synthetic oligonucleotides as primers to generate an SSM library ^52^. Importantly, SSM has been successfully applied to affinity maturation of antibodies, which are often screened in the scFv format because the presence of a single polypeptide chain eliminates difficulties with chain association that can occur with other antibody formats ^53^. Here, we generated a set of 266 site-directed variants of scFv735 in which all 19 amino acid substitutions were introduced at each of the 14 positions that contact polySia either directly or indirectly. Each SSM variant was individually expressed from plasmid pMAB in *E. coli* SHuffle T7 Express cells, collected in cell-free lysates, and screened for polySia-specific binding by enzyme-linked immunosorbent assay (ELISA) using chicken brain-derived polysialylated neural cell adhesion molecule (polySia-NCAM) as immobilized antigen. To confirm binding specificity, each SSM variant was also subjected to ELISA analysis using polySia-NCAM treated with endoneuraminidase N (endoN) which selectively cleaves linear polymers of sialic acid with α-2,8-linkage having a minimum length of 7-9 residues ^54^.

In the case of direct contact residues, Tyr-37, Arg-55, and Asp-232 were found to be intolerant to mutation, with substitutions to any other amino acid reducing binding to the polySia-NCAM antigen by ≥85% (**Fig. 2a and Supplementary Fig. 1**). Tyr-159 was nearly as intolerant to substitution, with binding activity reduced by ≥75% for all mutants except for the Y159R variant (scFv735^Y159R^), which preserved 75% of the binding measured for wild-type (wt) scFv735. The remaining two sites, Tyr-160 and Tyr-179, were relatively more tolerant. That is, while many substitutions at these positions abolished or dramatically reduced binding activity, several others exhibited binding that retained ≥50% of the binding measured for wt scFv735 and two – Y179W and Y179R – exhibited binding that exceeded that of wt scFv735. It should be noted that none of the variants exhibited any meaningful binding to an endoN-treated NCAM control antigen, confirming the polySia-specificity of all productive binders (**Supplementary Fig. 1**). Collectively, these SSM results confirm that all six direct contact residues are important for polySia binding, with four proving to be indispensable for this activity and two others exhibiting some degree of plasticity.

**Figure 2.**
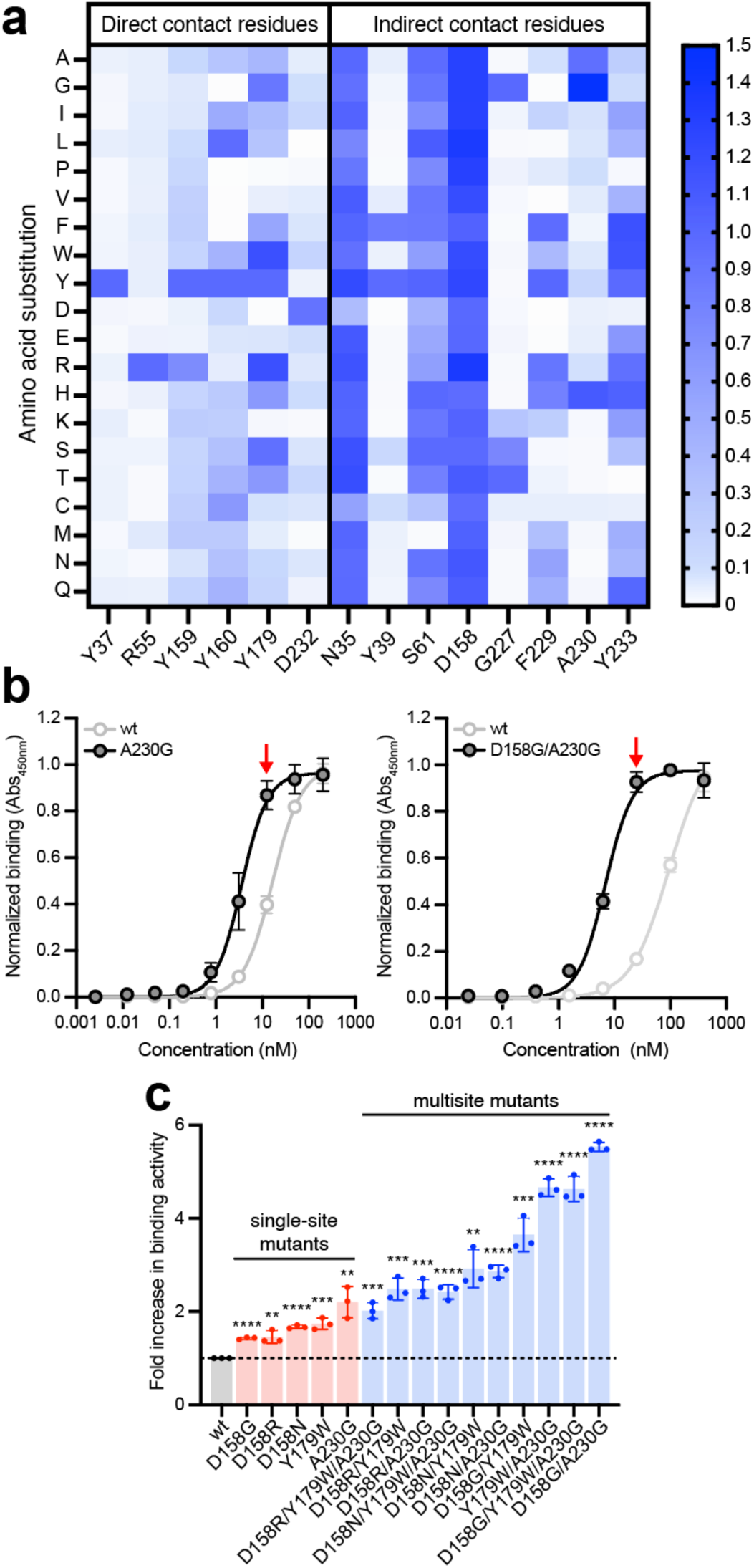
PolySia-specific binding of single- and multisite variants of scFv735. (a) Binding analysis of six direct contact residues (L:Tyr-37, L:Arg-55, H:Tyr-159, H:Tyr-160, H:Tyr-179, and H:Asp-232) and eight indirect contact residues (L:Asn-35, L:Tyr-39, L:Ser-61, H:Asp-158, H:Gly-227, H:Phe-229, H:Ala-230, and H:Tyr-233) in scFv735. Each contact residue was subjected to SSM, resulting in a set of 266 site-directed variants of scFv735 in which all 19 amino acid substitutions were introduced at each of the six positions. All 226 scFv735 variants and wt scFv735 were expressed from plasmid pMAB in *E. coli* SHuffle T7 Express cells and collected as cell lysates. Binding activity in cell-free lysates was quantified by ELISA using polySia- NCAM as immobilized antigen. An equivalent amount of total protein was loaded in each well. Heatmap data correspond to normalized ELISA signal for each variant relative to wt scFv735 (see **Supplementary** Figs. 1 and 2 for complete ELISA datasets). (b) Comparison of polySia binding activity for purified scFv735 variants harboring A230G and D158G/A230G substitutions (gray circles) versus wt scFv735 (white circles) as determined by ELISA with polySia-NCAM as immobilized antigen. Each variant was expressed from plasmid pMAB in *E. coli* strain SHuffle T7 Express cells and purified from cell-free lysate by Ni-NTA affinity chromatography. (b) Relative binding activity for select single-site and multisite variants as determined by ELISA with polySia-NCAM as immobilized antigen. Fold increase was calculated by normalizing the ELISA signal for each variant by the signal for wt scFv735 at an scFv concentration of 12.5 nM as denoted by the red arrows in (a). Data are average of biological replicates (*n* = 3) ± SD. Statistical significance was determined by unpaired two-tailed Student’s *t-*test. Calculated *p* values are represented as follows: **, *p* < 0.01; ***, *p* < 0.001; ****, *p* < 0.0001.

In the case of indirect contact residues, all eight newly tested sites were able to tolerate one or more substitutions without compromising binding whereas Arg-55, and Asp-232, which also doubled as direct contact sites, were completely intolerant to substitution (**Fig. 2a and Supplementary Fig. 2**). Asn-35, Ser-61, and Asp-158 exhibited the greatest plasticity, with nearly all substitutions (53 out of 57) conferring binding activity that was 50–125% of that measured for parental scFv735. In fact, every Asp-158 substitution showed binding activity equal to or greater than wt scFv735. In contrast, Tyr- 39 and a cluster of CDR H3 residues (Gly-227, Phe-229, Ala-230 and Tyr-233) were much less tolerant to mutation, with only a few substitutions at each of these sites leading to any significant activity. Interestingly, the A230G substitution conferred dramatically higher polySia-NCAM-specific binding relative to wt scFv735 even though all other Ala-230 substitutions except one (A230H) were completely inactive. Taken together, these results serve to highlight the important contribution that specific residues in CDR H3 make to antigen recognition.

### Identification of paratopic substitutions that confer enhanced binding affinity

Because a higher ELISA signal in cell-free extracts can result either from improved binding affinity or more efficient expression/folding, it was not possible to distinguish from the ELISA alone whether any SSM mutants exhibited higher affinity for polySia. Therefore, to more carefully quantify polySia binding, a total of 25 of the most active variants (marked by red asterisks in Supplementary Figs. 1 and 2) were purified from cell-free lysates and subjected to ELISA analysis with polySia-NCAM or endoN-treated NCAM to evaluate binding activity and specificity. Following this more rigorous sample preparation and normalization, the binding activity for many of the most active single-site mutants was found to be indistinguishable from wt scFv735, including several that initially showed enhanced activity compared to wt scFv735 in the non-purified, cell-free extracts (**Supplementary Fig. 3a**). At the same time, this analysis uncovered five variants all with substitutions in the V_H_ domain – scFv735^D158G^, scFv735^D158N^, scFv735^D158R^, scFv735^Y179W^, and scFv735^A230G^ – that exhibited stronger polySia-NCAM binding than wt scFv735, with binding activity of the mutants increasing by as much as ∼2.5-fold relative to wt scFv735 (**Fig. 2b-c and Supplementary Fig. 3b**). To determine if the elevated ELISA signals corresponded to enhanced binding affinity, we determined the intrinsic equilibrium dissociation constants (*K*_D_) for each of the single-site variants using biolayer interferometry (BLI). The measured *K*_D_ values for the five variants ranged from 23.2–44.0 nM while the *K*_D_ value for wt scFv735 was 72.7 nM (**Table 1 and Supplementary Fig. 4a**). The lower *K*_D_ values for the variants represented a ∼2–3-fold affinity enhancement, which was in good agreement with the ELISA results.

**Table 1.** Affinity of polySia-specific scFv and IgG variants

| Antibody clone | $K_D$ (nM) | $k_{on}$ ($10^4 \cdot M^{-1} \cdot s^{-1}$) | $k_{off}$ ($10^{-4} \cdot s^{-1}$) | $R^2$ value | Fold improved |
| --- | --- | --- | --- | --- | --- |
| *scFv735 variants\** |  |  |  |  |  |
| wt | 72.7 ± 0.3 | 1.09 ± 0.004 | 7.91 ± 0.02 | 0.990 | - |
| D158G | 23.2 ± 0.1 | 4.87 ± 0.024 | 11.3 ± 0.03 | 0.938 | 3.1 |
| D158N | 34.5 ± 0.1 | 2.33 ± 0.006 | 8.05 ± 0.02 | 0.986 | 2.1 |
| D158R | 44.0 ± 0.2 | 2.58 ± 0.010 | 11.3 ± 0.03 | 0.960 | 1.7 |
| Y179W | 39.4 ± 0.2 | 1.85 ± 0.006 | 7.30 ± 0.02 | 0.978 | 1.9 |
| A230G | 26.0 ± 0.1 | 1.90 ± 0.004 | 4.94 ± 0.01 | 0.992 | 2.8 |
| D158G/A230G | 20.4 ± 0.1 | 1.43 ± 0.002 | 2.93 ± 0.01 | 0.997 | 3.6 |
| H2Lib1-3 | 56.6 ± 0.2 | 1.81 ± 0.003 | 6.69 ± 0.02 | 0.984 | 1.3 |
| W177G | 41.5 ± 0.2 | 2.09 ± 0.007 | 8.69 ± 0.02 | 0.992 | 1.8 |
| D158N/W177G | 19.6 ± 0.1 | 7.22 ± 0.039 | 14.2 ± 0.04 | 0.995 | 3.7 |
| *chAb735 variants\** |  |  |  |  |  |
| wt | 15.7 ± 0.1 | 6.65 ± 0.025 | 10.4 ± 0.02 | 0.976 | - |
| D158G | 2.62 ± 0.01 | 15.1 ± 0.042 | 3.96 ± 0.01 | 0.976 | 6.0 |
| A230G | 5.43 ± 0.02 | 8.54 ± 0.022 | 4.64 ± 0.01 | 0.987 | 2.9 |
| D158G/A230G | 2.28 ± 0.01 | 9.02 ± 0.015 | 2.05 ± 0.01 | 0.994 | 6.8 |
| H2Lib1-3 | 10.9 ± 0.06 | 9.77 ± 0.046 | 10.7 ± 0.03 | 0.965 | 1.4 |
| W177G | 9.09 ± 0.04 | 7.56 ± 0.025 | 6.87 ± 0.02 | 0.986 | 1.7 |
| D158N/W177G | 2.82 ± 0.01 | 9.00 ± 0.015 | 2.53 ± 0.01 | 0.995 | 5.6 |
| *chAb735 variants (on-cell $K_D$)\*\** |  |  |  |  |  |
| wt | 55.7 | - | - | 0.972 | - |
| D158G | 6.80 | - | - | 0.918 | 8.2 |
| A230G | 16.2 | - | - | 0.967 | 3.4 |
| D158G/A230G | 6.10 | - | - | 0.971 | 9.1 |
| H2Lib1-3 | 24.4 | - | - | 0.987 | 2.3 |
| W177G | 16.7 | - | - | 0.989 | 3.3 |
| D158N/W177G | 16.3 | - | - | 0.974 | 3.4 |

\*Binding parameters were fitted according to the Langmuir model in Octet Analysis Studio.
\*\*On-cell apparent affinity was determined by non-linear regression analysis in Prism 10.
Note: the H2Lib1-3 variants carried four CDR H2 mutations: W177R, I178V, Y179F, and G181K.

### Creation of higher affinity binders from simple combinatorial mutagenesis

When several affinity enhancing mutations are discovered as separate hits in a single round of mutagenesis and screening, combining these mutations can be beneficial. Therefore, to determine whether combination of the five single-site mutations could yield even higher affinity variants, we constructed a small combinatorial beneficial mutation (CBM) library containing all two- and three-site permutations of the individual beneficial mutations. The resulting ten variants were individually expressed and purified from *E. coli* SHuffle T7 Express cells and subsequently screened by ELISA. Whereas CBM variants containing either the D158N or D158R substitution showed only modest improvement relative to their single-site mutant counterparts, those involving D158G were greatly improved, with increases of ∼2–3 fold above the corresponding single-site variants and ∼4–6 fold above wt scFv735 (**Fig. 2b-c**). In line with these improvements, BLI analysis of the most active CBM variant, scFv735^D158G/A230G^, revealed significant affinity enhancement with a measured *K*_D_ value of 20.4 nM (**Table 1 and Supplementary Fig. 4a**).

### Isolation of affinity-matured scFv735 variants from CDR-focused ‘NNK’ libraries

Although rationally introducing mutations into contact residues has theoretical advantages, we considered that mutations to other CDR residues not in contact with the antigen could lead to improved affinity. Indeed, mutagenesis studies often yield substituted CDR residues that are not in contact with the antigen ^55^. Therefore, to identify additional residues within the CDRs that might enhance affinity, we pursued a semi- rational approach whereby the V_H_-domain CDRs in scFv735 were subjected to saturation mutagenesis using degenerate NNK primers ^56^ and the resulting CDR-focused NNK libraries were screened by yeast surface display. We focused exclusively on the V_H_ CDRs because CDR-L3 makes no direct contacts with polySia and because all mutations identified in our SSM/CBM screen that conferred improved binding were in the V_H_ domain. Accordingly, we constructed six independent, six-codon NNK libraries that collectively spanned the three V_H_ CDRs and flanking residues (**Fig. 3a**), which were each fused in- frame to the gene encoding the yeast Aga2p protein in the yeast surface display plasmid pCT-CON ^57^ using homologous recombination. We restricted the complexity of each NNK library to six residues because the number of sequences in a library of more than six mutated residues would exceed the practical limits of yeast display. The integrity and diversity of each library was assessed by sequencing ∼25 random clones. In general, the libraries were highly diverse with each containing ∼10^8^ unique clones and an extremely low number of non-mutated wt clones.

**Figure 3.**
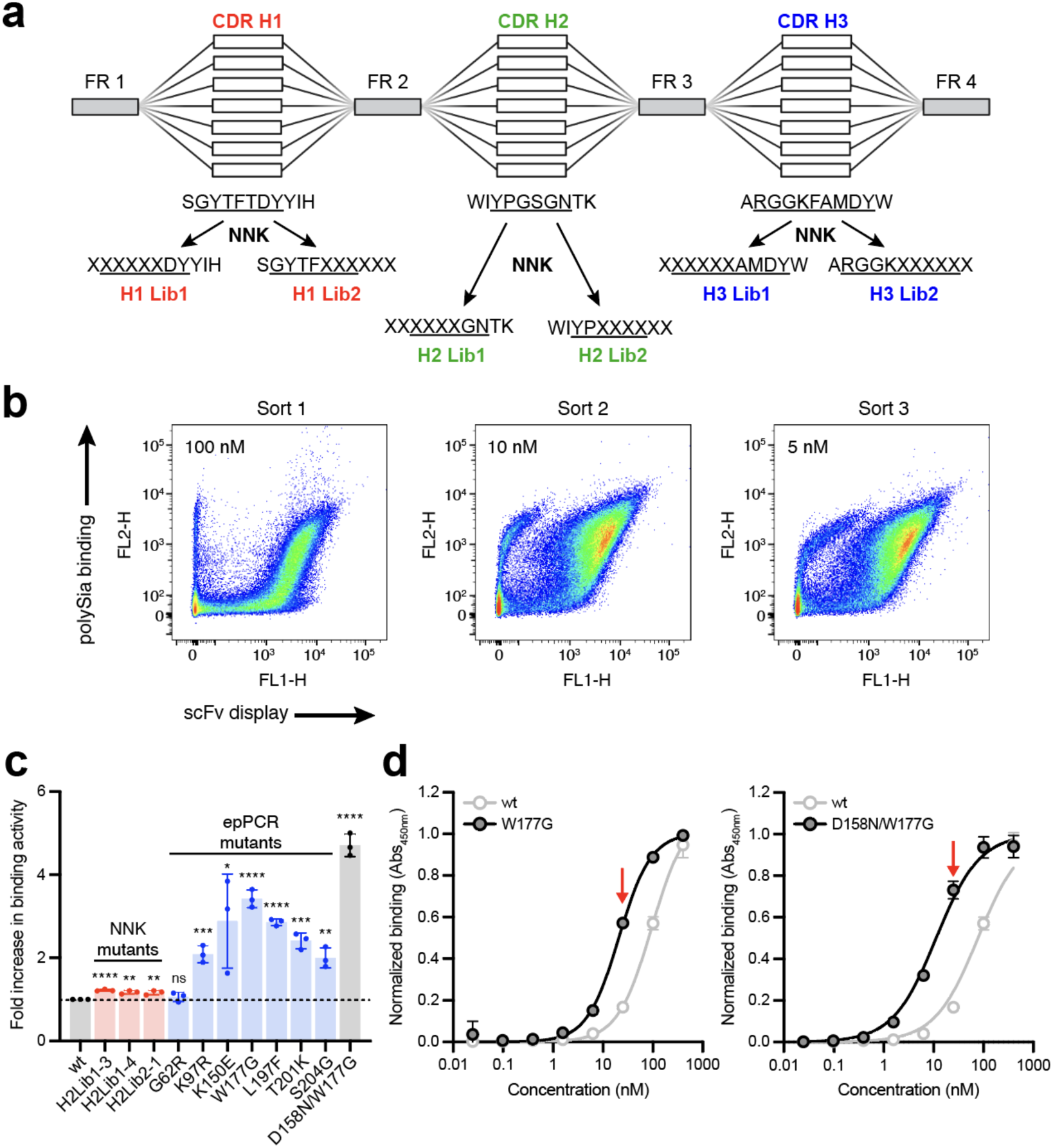
Discovery of affinity-matured scFv735 variants from combinatorial library screening. (a) Schematic of NNK library construction leading to six independent, six-codon NNK libraries that collectively spanned the three VH CDRs (underlined text) and flanking residues of scFv735. (b) Representative scatter plots corresponding to FACS-based screening of yeast display libraries. Yeast libraries were labeled with rabbit anti-Myc tag antibody followed by Alexa Fluor 647-conjugated goat anti-rabbit IgG antibody for scFv display and biotinylated polySia-NCAM followed by streptavidin-Alexa Fluor 488 conjugate for polySia binding. Yeast cells were stained with decreasing concentrations of biotinylated polySia-NCAM (100, 10, and 5 nM) over three rounds of FACS. The top 0.1-0.3% cells were selected from the sort gates (see **Supplementary Fig. 7** for gating strategy). (c) Relative binding activity for NNK- and epPCR-derived hits as determined by ELISA using polySia-NCAM as immobilized antigen. Fold increase was calculated by normalizing the ELISA signal for each variant by the signal for wt scFv735 at an scFv concentration of 25 nM as denoted by red arrows in (d). Statistical significance was determined by unpaired two-tailed Student’s *t-*test. Calculated *p* values are represented as follows: *, *p* < 0.05; **, *p* < 0.01; ***, *p* < 0.001; ****, *p* < 0.0001; ns, not significant. (d) Comparison of polySia binding activity for purified scFv735 variants harboring W177G and D158N/W177G substitutions (gray circles) versus wt scFv735 (white circles) as determined by ELISA using polySia-NCAM as immobilized antigen. Each variant was expressed from plasmid pMAB in *E. coli* strain SHuffle T7 Express cells and purified from lysate by Ni-NTA affinity chromatography. Data are the average of biological replicates (*n* = 3) ± SD.

The resulting plasmid libraries were used to transform *Saccharomyces cerevisiae* strain EBY100, which enables cell-surface expression of recombinant scFv antibody libraries ^58^. Following several rounds of negative selection using magnetic-activated cell sorting (MACS) to deplete the libraries of binders to undesired targets (e.g., endoN- treated NCAM), an additional round of MACS was performed to positively select binders to polySia-NCAM. Next, four rounds of fluorescence-activated cell sorting (FACS) were performed with polySia-NCAM at successively decreasing concentrations (100 nM, 10 nM, and 5 nM) (**Fig. 3b**). During the final round of FACS sorting, individual yeast cells representing putative polySia-positive clones were sorted into 96-well plates, after which plasmid DNA was isolated and sequenced. All unique sequences were subcloned into plasmid pMAB to enable *E. coli*-based expression and purification of scFv735 variants, which were subsequently analyzed by ELISA. From the six NNK libraries, we isolated 3 unique clones, all from CDR H2-focused libraries, that displayed only modestly improved polySia-NCAM binding activity relative to wt scFv735 (**Fig. 3c and Supplementary Fig. 5**). Among these, scFv735^H2Lib1–3^ was the most significantly improved clone with a 1.2- fold increase in binding activity (**Fig. 3c**) that translated to a similarly modest affinity enhancement (**Table 1**). This clone carried a total of four mutations, two within CDR H2 (Y179F, G181K) and two just upstream (W177R, I178V), with the Trp-177 and Gly-181 residues being mutated in all three NNK hits (**Supplementary Fig. 5**).

### Isolation of affinity-matured scFv735 variants from error-prone PCR library

Having explored rational and semi-rational mutagenesis strategies, we next investigated random mutagenesis of the entire variable region of scFv735 using error-prone PCR (epPCR) ^59^. This decision was motivated in part by our isolation of mutations outside the CDR that appeared to contribute to enhanced affinity. We were further motivated by earlier observations that framework region (FR) mutations can indirectly impact antibody-antigen interactions by affecting the antibody conformation ^60^ or by the positioning of contact residue side chains ^61^. FR mutations have even been found to alter the electrostatic surface potential of the antigen binding site of anti-glycan antibody, increasing its positive charge in a manner that enhances long-range interactions with its negatively charged carbohydrate antigen ^35^. During the epPCR mutagenesis process, we targeted a mutational frequency of ∼0.5% (<4 random substitutions per 735-bp scFv735 gene) so that library members would harbor a similar number of mutations as our SSM/CBM- and NNK-derived variants. We screened the epPCR library following an identical yeast surface display procedure as above and identified 7 unique clones, all of which were single-site substitutions that displayed improved polySia-NCAM binding activity relative to wt scFv735 (**Fig. 3c and Supplementary Fig. 6**). To our surprise, all isolated mutations occurred in FRs outside the CDRs, although several (G62R, K79R, K150E, and W177G) were located near CDRs and/or contact residues. For example, affinity-matured scFv735^W177G^ harbored a W177G substitution, which is just upstream of an important direct contact residue, Tyr-179, in CDR H2. This variant was the most improved epPCR- derived clone with a nearly 3.5-fold increase in both polySia-NCAM binding activity (**Fig. 3c-d**) and affinity (**Table 1**).

The fact that Trp-177 was also mutated independently in the three NNK-derived variants suggested a critical role for this residue in high-affinity polySia binding. Therefore, we performed combinatorial mutagenesis to combine the W177G mutation with the best single-site substitutions from the SSM analysis, namely D158G/N/R, Y179W, and A230G. After screening all pairwise combinations, one clone in particular, scFv735^D158N/W177G^, exhibited greatly improved polySia-NCAM binding activity along with substantially enhanced affinity (**Fig. 3c-d** and **Table 1**). In fact, the improvement seen for scFv735^D158N/W177G^ rivaled that of scFv735^D158G/A230G^, with measured *K*_D_ values of 19.6 and 20.4 nM, respectively, that were 3-fold improved over wt.

### Reformatting scFvs into full-length IgGs yields low nanomolar polySia binders

Because full-length monoclonal antibodies (mAbs) are often the preferable format for many follow-on applications, we proceeded to convert a subset of our best affinity- matured scFv735 variants into full-length chimeric antibody (chAb) variants. The reformatting process involved cloning the V_H_ and V_L_ genes of each scFv into the plasmid pVITRO1, which contains the constant region (Fc) of human IgG1 ^62^. The pVITRO1- encoded IgG1 variants were produced as secreted IgGs from stably transfected Freestyle 293-F cells in 125-mL spinner flasks. After protein A affinity column purification, all chAb735 variants were over 95% pure as judged by reduced SDS-PAGE, with yields in the ∼5-10 mg/L range (**Supplementary Fig. 8a**).

Each of the purified chAb735 variants exhibited improved polySia-NCAM binding activity relative to parental chAb735 (**Fig. 4a and Supplementary Fig. 8b**), confirming that the binding improvements were generally retained following reformatting. Likewise, all chAb735 variants exhibited higher affinity than parental chAb735, with *K*_D_ values for the three best affinity-matured variants – chAb735^D158G^, chAb735^D158G/A230G^, and chAb735^D158G/W177G^ – reaching 2.62, 2.28, and 2.82 nM, respectively, compared with 15.7 nM for parental chAb735 (**Table 1 and Supplementary Fig. 4b**). The 5-7-fold increases in affinity observed for these three chAb735 variants were even greater than the enhancement conferred by the same substitutions in the scFv format. It should be noted that the *K*_D_ measured for wt chAb735, 15.7 nM, was in good agreement with the affinity values reported previously for mAbs composed of the same variable regions ^63, 64^. It is also worth noting that the *K*_D_ value for wt chAb735 was nearly 5-fold stronger than that measured for wt scFv735, consistent with the frequently observed phenomenon of mAbs having higher affinities than their scFv counterparts ^65^ including in the context of antibodies that bind carbohydrate antigens ^35^.

**Figure 4.**
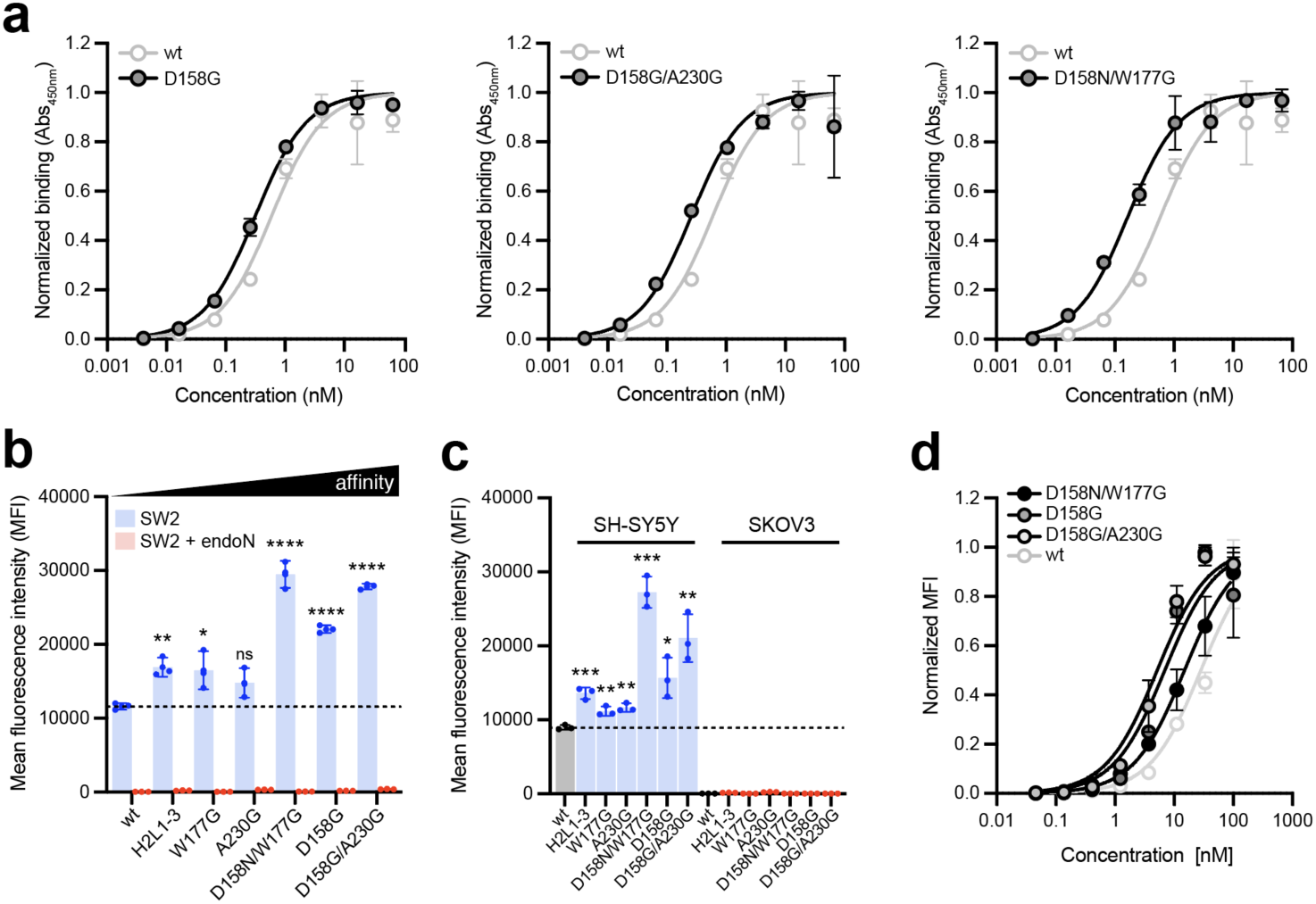
PolySia-specific binding analysis of reformatted chAb735 variants. (a) Comparison of polySia binding activity for purified chAb735 variants harboring D158G, D158G/A230G and D158N/W177G substitutions (gray circles) versus wt chAb735 (white circles) as determined by ELISA. Each variant was expressed from plasmid pVITRO1 in Freestyle 293-F cells and purified from culture supernatants by protein A affinity chromatography. PolySia-NCAM was used as the immobilized antigen and data are the average of biological replicates (*n* = 3) ± SD. (b) Binding of chAb735 constructs to SW2 tumor cells with high levels of polySia expression (blue) or without polySia expression due to endoN treatment (red). A total of 10^6^ SW2 cells were stained with 33.3 nM of indicated chAb735 variant, followed by staining with Alexa Fluor 488- conjugated goat anti-human IgG secondary Ab. Mean fluorescence intensity (MFI) data are the average of biological replicates (*n* = 4) ± SD. Statistical significance was determined by unpaired two-tailed Student’s *t-*test. Calculated *p* values are represented as follows: *, *p* < 0.05; **, *p* < 0.01; ***, *p* < 0.001; ****, *p* < 0.0001; ns, not significant. (c) Same as in (b) but with SH-SY5Y (blue) or SKOV3 (red) cells. (d) Retention of chAb735 constructs on SW2 cells. Cells were incubated with serially diluted chAb constructs (1:3 starting from 100 nM) at room temperature for 30 min after which chAbs retained on cell surface after washing were detected by Alexa Fluor 488-conjugated goat anti-human IgG Ab. MFI data represent the amount of IgGs on the cell surface and are the average of biological replicates (*n* = 3) ± SD.

### Affinity-matured chAb735 variants efficiently opsonize polySia-expressing cells

To further evaluate the functional consequences of affinity maturation, we next investigated the ability of the chAb735 variants to bind polySia in its native context. To this end, we assessed binding specificity using a flow cytometric binding assay with the human small-cell lung carcinoma (SCLC) cell line SW2, which is highly polySia-positive^63^. Following incubation with SW2 cells, clear cell surface staining was detected for all chAbs, with the highest affinity variants binding most strongly as evidenced by the 2-3- fold increase in staining intensity (**Fig. 4b**). Enhanced cell staining by higher affinity antibodies is a phenomenon that has been observed previously, for example, with affinity- modulated anti-EGFR antibodies ^66, 67^. In contrast, no cell-surface staining was detected for any of the antibodies when the SW2 cells were pretreated with endoN, which selectively removes polySia from the cell surface ^63^. Given the diversity of glycans found on SCLC tumor cells ^68, 69^, including monosialylated and disialylated structures with linkages other than α2,8, the lack of binding to endoN-treated SW2 cells provided strong evidence that the exquisite polySia specificity seen previously ^63^ was retained by the affinity-matured variants. This polySia-specific binding was significant because affinity maturation of anti-glycan antibodies is usually accompanied with epitope spread, resulting in loss of specificity ^33, 36^. To further confirm polySia specificity, we investigated cell surface binding using two additional human cell lines, the polySia-positive neuroblastoma cell line SH-SY5Y and the polySia-negative ovarian cancer cell line SKOV3. In support of the strict polySia specificity of these antibodies, we observed robust staining of SH-SY5Y but not SKOV3 cells following incubation with each of the chAbs (**Fig. 4c**). As with SW2 cells, the most intense staining of SH-SY5Y cells was observed for the highest affinity chAb735 variants, with relative staining levels across these two cell lines nearly indistinguishable.

To better quantify the differences in cell binding, we measured the apparent affinity of each chAb using our on-cell binding assay as described ^70, 71^. Apparent affinity is an important determinant of IgG binding to polySia-expressing tumor cells. When the chAb735 variants were tested for binding to SW2 cells at a variety of concentrations, the three highest affinity clones exhibited roughly equivalent tumor cell retention that were greater than that achieved using parental chAb735 (**Fig. 4d**), with apparent *K*_D_ values in the 6.1-16.7 nM range for the chAb735 variants compared to 55.7 nM for wt chAb735 (**Table 1**). Although these apparent affinities were generally 2-3-fold lower than the intrinsic affinities calculated from BLI analysis, the fold improvements for each variant relative to parental chAb735 were similar, providing further evidence of affinity maturation.

### Higher affinity chAb735 variants exhibit superior tumor cell killing

Having confirmed the cellular binding properties of the chAb735 variants by flow cytometry, we next evaluated their ADCC and CDC activity, which together provide a comprehensive understanding of immune-mediated cytotoxic mechanisms. We speculated that affinity-optimized chAb735 variants would trigger greater ADCC and CDC owing to their more pronounced opsonization of polySia-positive tumor cells. Indeed, the affinity-matured chAb735 variants mediated superior ADCC relative to parental chAb735 as determined using human SCLC SW2 cells as target in the presence of human peripheral blood mononuclear cells (PBMCs) as effectors (**Fig. 5a**). These same chAb735 variants also promoted superior CDC relative to parental chAb735 as determined using target SW2 cells in the presence of baby rabbit complement (**Fig. 5b**). As expected, little to no cytotoxicity was observed in assays performed using an isotype control chAb or using anti-polySia chAbs but in the absence of effector PBMCs or complement. The strongest ADCC and CDC activities were observed for the highest affinity variant, chAb735^D158G/A230G^ (*K*_D_ ≈ 2.28 nM), which promoted significantly greater cytotoxicity against SW2 cells relative to wt chAb735 (*K*_D_ ≈ 15.7 nM) at all antibody concentrations tested (**Fig. 5a-b**).

**Figure 5.**
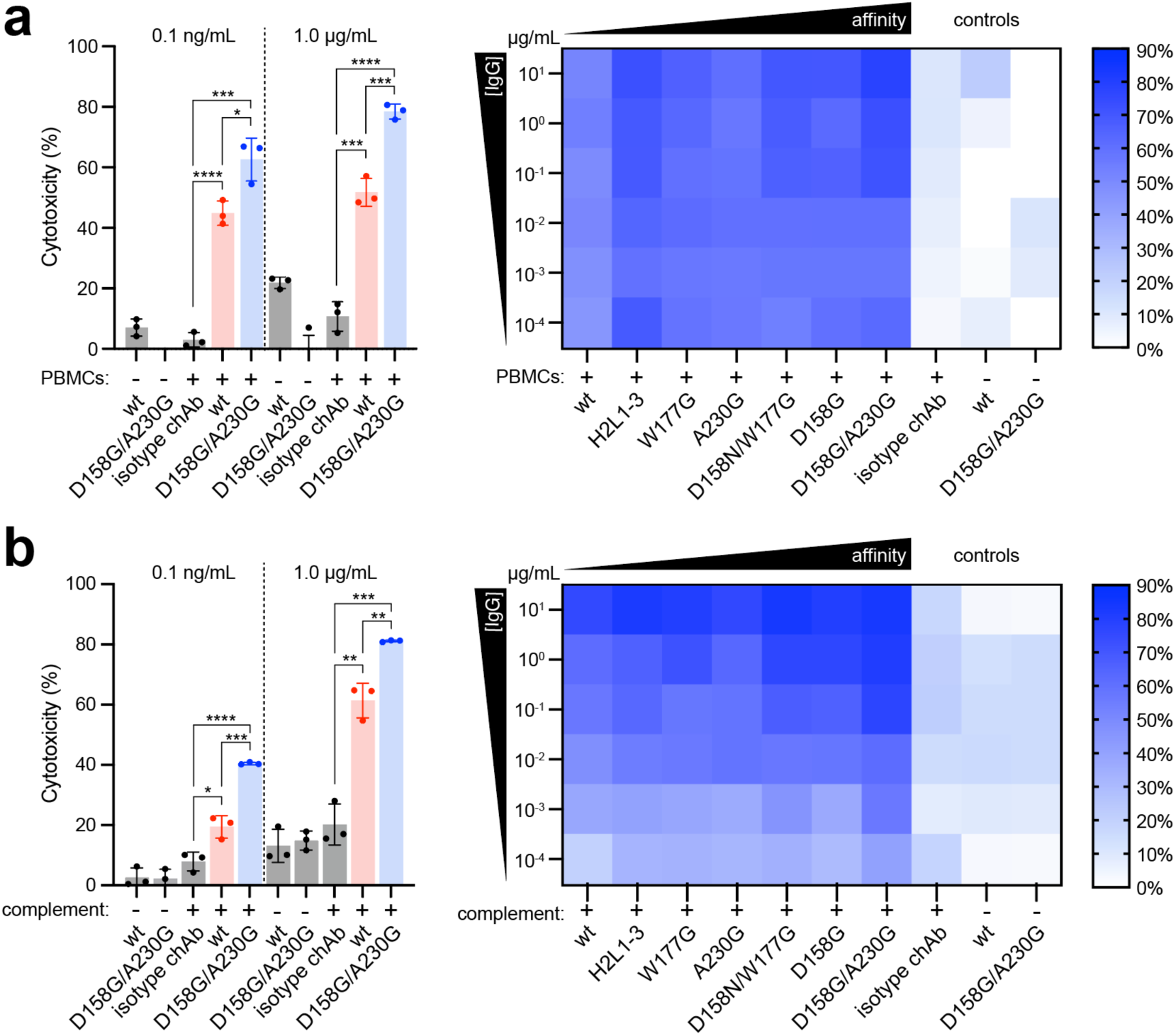
Cytotoxicity of chAb735 variants against polySia-expressing tumor cells. (a) ADCC and (b) CDC activity of chAb735 variants (blue bars) and wt chAb735 (red bars) against SW2 tumor cells. For ADCC assays, effector PBMCs to target SW2 cell ratios of 20:1 were used. For CDC assays, baby rabbit complement was used at a final concentration of 10% (v/v). Cytotoxicity was assayed over a range of IgG concentrations in the presence (+) or absence (-) of PBMCs or complement, with representative results depicted for 0.1 ng/mL and 1 μg/mL in bar graphs at left. Isotype chAb in the presence of PBMCs/complement or chAb735 and chAb735^D158G/A230G^ in the absence of PBMCs/complement served as negative controls (gray bars). Data are average of biological replicates (*n* = 3) ± SD. Statistical significance was determined by unpaired two-tailed Student’s *t-*test. Calculated *p* values are represented as follows: *, *p* < 0.05; **, *p* < 0.01; ***, *p* < 0.001; ****, *p* < 0.0001; ns, not significant.

## DISCUSSION

In light of the important role that carbohydrate antigens play in cancer biology and infectious disease, antibodies targeting such antigens have enormous potential for both basic research and clinical applications ^10^. However, carbohydrates remain an underexplored target space for mAb development. Α major impediment to forward progress is the fact that carbohydrate molecules are typically T cell-independent antigens, leading to weaker and less robust immune responses and the elicitation of primarily IgM antibodies that are low affinity and broadly specific ^19–21^. Indeed, early efforts to induce antibodies to polySia using tetanus toxoid (TT) or keyhole limpet hemocyanin (KLH) conjugates failed to elicit significant immune responses in immunized mice, rabbits, and humans ^72, 73^. Likewise, immunization of wildtype BALB/c mice with *Neisseria meningitidis* group B, which express cell surface polySia, only triggered IgM but not IgG antibodies specific for polySia ^38^. The poor immunogenicity of polySia has been attributed to immunologic tolerance that arises due to its resemblance with structures present in mammalian hosts (i.e., self-antigens). Yet even after chemical alteration of the polySia structure to render it more immunogenic, conjugates bearing this carbohydrate elicited IgM but not IgG antibodies against polySia ^73^. To date, the only successful efforts to elicit class-switched IgGs with polySia specificity required systems capable of immunological hyperreactivity, i.e., immunizing autoimmune NZB mice with *N. meningitidis* group B ^38^ or immunizing wildtype BALB/c mice with strongly immunostimulatory outer membrane vesicles (OMVs) engineered to display polySia antigens on their exterior ^74^. Of relevance to the studies herein, the NZB system was used to generate an IgG2a monoclonal antibody against polySia, namely mAb735. However, like other anti-glycan mAbs that have been derived from animal immunization, mAb735 is essentially the germline sequence (V_L_ differing from IGKV1-110*02 by just one FR2 mutation and V_H_ differing from IGHV1-84*01 by just 3 FR mutations, as inferred from germline genes identified via IgBlast ^75^) and exhibits modest affinity (*K*_D_ ≈ 15.7 nM for the chimeric version of mAb735 as measured here) ^43, 46^.

While affinity improvement via protein engineering is an attractive approach to overcome the suboptimal binding properties of anti-glycan mAbs and potentially increase their therapeutic index, only a handful of efforts have been reported so far ^27, 30–37^. Therefore, to expand the target space, in the current study we applied both rational and random molecular evolution techniques to identify mutations in the variable regions of mAb735 that increased its polySia-binding activity. For the rationally designed variants, we explored structure-guided SSM of key residues that were identified based on an available crystal structure of scFv735 in complex with octasialic acid ^45^. Our success in isolating higher affinity variants using SSM illuminates how the maturation process can be accelerated by deducing the contact residues from antibody-antigen complex structures when available. In parallel, we also exploited semi-random (i.e., CDR mutagenesis via NNK libraries) and entirely random (i.e., epPCR of the entire Fv) mutagenesis strategies, in which stepwise decreases in polySia antigen concentration during FACS resulted in the enrichment of high-affinity binders. Although mutating sites within CDR loops, either by rational SSM or semi-random NNK mutagenesis, has theoretical advantages over mutagenesis of the entire variable regions, it is not uncommon for FR mutations to exert beneficial effects on antigen binding, for example, by affecting the antibody conformation ^60^, re-positioning contact residue side chains ^61^, or altering electrostatic surface potential ^35^. Indeed, all higher affinity variants that we isolated from the epPCR library carried just a single FR mutation. That said, the greatest affinity improvements were associated with CDR mutations. Among these, the combined D158G/A230G substitutions increased polySia-specific affinity of chAb735—a chimeric version of mAb735—by nearly 7-fold (*K*_D_ ≈ 2.28 nM). This enhanced affinity translated into a ∼2-fold improvement in both ADCC and CDC, in line with other studies showing that higher affinity antibodies promote more effective *in vitro* tumor cytotoxicity against both protein ^76^ and glycan antigens ^30, 35^.

In summary, by using a combination of rational design and directed evolution, we were able to engineer affinity-maturated antibodies with single or double mutations that conferred significant improvements in polySia-binding affinity and *in vitro* tumor cell opsonization and cytotoxicity, while maintaining restricted polySia-positive tumor cell cross-reactivity. The identification of important affinity enhancing residues in both the CDRs and FRs of mAb735 not only provides greater insight into the molecular mechanisms by which polySia antigen is specifically recognized by antibodies but also furnishes improved clinical candidates against an important carbohydrate tumor antigen^44^. Moreover, the effective affinity maturation approaches described here offer a roadmap for empowering the functional enhancement of other anti-glycan mAbs in the future, with an eye towards creating next-generation molecules with higher therapeutic potential.

## MATERIALS AND METHODS

### Strains and plasmids

*E. coli* strain DH5α was used for all plasmid construction. *E. coli* strain SHuffle T7 Express (New England Biolabs) ^77^ was used for scFv expression and purification as well as library selections. *S. cerevisiae* strain EBY100 was used for yeast surface display. Plasmid pMAB was used for *E. coli-*based expression and purification of all scFvs. Plasmid pCT-CON ^57^ was used for expression of all scFv libraries in yeast and was kindly provided by Dr. Dane Wittrup (MIT). SDM of select scFv735 residues, either individually or in combination, was performed by amplifying the entire pMAB-scFv735- 6xHis plasmid by inverse PCR using primers carrying specific mutations or degenerate NNK primers followed by transformation of chemically competent DH5α cells with the resulting linear PCR products. Plasmid pVITRO-735-IgG1/κ ^63^, which was constructed previously from pVITRO1-Trastuzumab-IgG1/κ (Addgene plasmid #61883) ^62^, was used for expression of wt chAb735. All chAb735 derivatives were constructed by subjecting plasmid pVITRO-735-IgG1/κ to SDM according to standard protocols. All plasmid DNA was isolated from overnight cultures using a QIAprep Spin Miniprep Kit (QIAgen) and subsequently confirmed by Sanger sequencing at the Genomics Core Facility of the Cornell Biotechnology Resource Center (BRC).

### Mammalian cell culture

Freestyle 293-F cells (ThermoFisher) were used for the expression of all full-length chAbs from pVITRO1-based plasmids. Freestyle 293-F cells were maintained in FreeStyle 293 expression medium (ThermoFisher). SH-SY5Y (cat # CRL-2266) and SKOV3 (cat # HTB-77) cell lines were obtained from ATCC and SW2 cells were kindly provided by Dr. Karen Colley (University of Illinois at Chicago). SH-SY5Y cells were maintained in high-glucose DMEM/F12 medium supplemented with 10% (v/v) fetal bovine serum (FBS) (VWR), 1% (v/v) MEM nonessential amino acids solution (ThermoFisher), penicillin (100 U/mL), and streptomycin (100 μg/mL) (ThermoFisher). SKOV3 and SW2 cells were cultured in high-glucose DMEM supplemented with 10% (v/v) FBS, penicillin (100 U/mL) and streptomycin (100 μg/mL). All cell lines are maintained at 37 °C in a 5% CO_2_-humidified incubator.

### Library construction

Overlap extension PCR was used for the construction of scFv sequences harboring CDR-focused ‘NNK’ libraries. Overlapping forward and reverse primers containing NNK degenerate codons were designed to specifically mutate six consecutive amino acids within each mutated CDR region while a second pair of overlapping primers was designed to bind elsewhere in the pCT-CON backbone. PCR amplification using these primers and plasmid pCT-CON-scFv735 as template was performed, yielding two fragments that were subsequently assembled by overlap PCR to create a complete scFv product. For the construction of the epPCR library, the entire scFv735 sequence was randomly mutated by PCR with a GeneMorph II random mutagenesis kit (Stratagene) according to the manufacturer’s instructions. The mutation frequency was designed to introduce 0-9 base pair mutations per kb. The final scFv products from these reactions, which represented NNK and epPCR libraries, were isolated using a QIAquick PCR Purification Kit (Qiagen). All scFv libraries and linearized pCT-CON were introduced into EBY100 cells by electroporation, resulting in yeast cell libraries consisting of ∼10^8^ members.

### Selection of scFv735 mutants from yeast surface display libraries

Yeast libraries were grown in 200 mL SD-CAA media (20 g/L D-glucose, 6.7 g/L yeast nitrogen base, 5 g/L casamino acids, 5.4 g/L Na_2_HPO_4_, and 8.6 g/K NaH_2_PO_4_) overnight at 30 °C. The day before each sort, an amount of cells totaling 30x the diversity of the library was induced for scFv expression and surface display by switching the media to SG-CAA (18 g/L galactose, 2 g/L D-glucose, 6.7 g/L yeast nitrogen base, 5 g/L casamino acids, 5.4 g/L Na_2_HPO_4_, and 8.6 g/K NaH_2_PO_4_) and incubating overnight at 20 °C. The target antigen, polysia-NCAM (Sigma-Aldrich) was biotinylated using the EZ-Link-Sulfo-NHS-LC- Biotinylation kit (ThermoFisher). Biotinylated polysia-NCAM (33 pM) was incubated with 4 x 10^6^ streptavidin beads (ThermoFisher) in 100 μL PBSA (PBS with 0.1% (w/v) BSA) for 2 h at 4 °C. Before FACS selection, induced yeast library was incubated with the beads coated with biotinylated polysia-NCAM for 2 h at 4 °C, followed by the separation with a magnetic stand. The isolated beads were washed three times with PBSA, added to 5 mL SD-CAA media, and grown overnight in a shaking incubator at 30 °C. The recovered yeast cells were induced in SG-CAA media overnight at 20 °C. Approximately 5×10^7^ yeast cells were pelleted, washed twice with PBSA, and resuspended first in 100 nM biotinylated polysia-NCAM and a 1:100 dilution of rabbit anti-Myc tag antibody (ThermoFisher). After incubation, yeast cells were washed three times and then resuspended in 200 μL of PBSA buffer. 1:100 dilutions of both Alexa Fluor 647- conjugated goat anti-rabbit IgG antibody (ThermoFisher) and Streptavidin Alexa Fluor 488 Conjugate (ThermoFisher) were added, incubated at 4 °C for 30 min, and washed three times with PBSA buffer prior to FACS-based selection. Cells were sorted using a BD FACSMelody Cell Sorter (BD Biosciences). Sorting gates were set to acquire 0.1% of the population with the highest binding signal. Cells were sorted into 1 mL SD-CAA and were grown overnight at 30 °C. Cells were then induced in SG-CAA for the next round of sorting. For the next two selections, ∼1×10^7^ yeast cells were used for staining with biotinylated polysia-NCAM, and the antigen concentration was decreased to 10 nM and 5 nM in the consecutive rounds. Plasmids were isolated from yeast using the Zymoprep Yeast Plasmid Miniprep II Kit (Zymo Research) according to the manufacturer’s instructions. Following transformation of chemically competent DH5α by heat shock, all yeast-derived plasmids were isolated from overnight cultures using a QIAprep Spin Miniprep Kit (QIAgen), after which sequences were confirmed by Sanger sequencing at the Genomics Core Facility of the Cornell BRC.

### Expression and purification of scFv antibodies

For scFv expression, sequenced plasmids were used to transform chemically competent SHuffle T7 Express cells, after which a single colony was used to inoculate 5 mL LB supplemented with 100 μg/mL carbenicillin or ampicillin, and grown overnight at 37 °C. The next day, 5 mL fresh LB supplemented with 100 μg/mL carbenicillin or ampicillin was inoculated 1/100 with the overnight culture and cells were grown at 37 °C until reaching a density corresponding to an absorbance measured at 600 nm (Abs_600_) ≈ 0.5–0.8. At this point, scFv expression was induced by addition of 0.1 mM IPTG, after which cells were incubated an additional 16 h at 30 °C. Cells were harvested by centrifugation (16,000 ×g, 4 °C) and resuspended in 1 mL BugBuster 10x Protein Extraction Reagent (Millipore). To prevent protein degradation, 10 μL of Halt Protease Inhibitor Cocktail (100x) (ThermFisher) was added. After incubation for 30 min, cell lysate was collected by centrifugation at 16,000 ×g for 20 min at 4 °C. Total protein in cell lysates was quantified by Bradford assay.

For scFv purification, cells were grown and induced identically as above, after which harvested cells were resuspended in PBS and lysed using an EmulsiFlex-C5 homogenizer (Avestin). The cell lysate was clarified by centrifugation at 15,000 ×g at 4°C for 25 min and then mixed with HisPur Ni-NTA Resin (ThermoFisher). The resin-lysate mixture was incubated at room temperature with end-over-end mixing for 30 min. The mixture was then applied to a polypropylene gravity column and the lysate was allowed to completely pass through the column. The resin was then washed with 3x column volumes of wash buffer containing PBS supplemented with 25 mM imidazole and the protein was eluted with PBS supplemented with 250 mM imidazole. Purified fractions were applied to protein concentrators (10K MWCO; ThermoFisher) to change the buffer to PBS.

### Expression and purification of full-length IgG antibodies

IgGs were expressed in Freestyle 293-F suspension cells as described previously ^62^. Briefly, FreeStyle 293-F cells were grown in FreeStyle 293 Expression Medium (ThermoFisher). pVITRO1 plasmids encoding chAb735 and its variants were isolated from DH5α using QIAprep Spin Midiprep Kit (QIAgen) followed by transfection into FreeStyle 293-F cells using FreeStyle MAX transfection reagent (ThermoFisher) according to the manufacturer’s instructions and selected with hygromycin B. Culture media was initially collected every 72 h, followed by every 48 h after selection. Collected culture media was centrifuged at 1,000 ×g for 15 min, passed over 0.2-μm filters (VWR), and stored at 4 °C until use. Protein A agarose resin (Mabselect SuRe) was used to purify antibodies from the supernatant according to the manufacturer’s instructions. The resin was equilibrated with PBS in a polypropylene gravity column. The supernatant was allowed to completely pass through the column. The resin was then washed with PBS and IgGs were eluted from the column with 0.1 M glycine-HCl (pH 3) and neutralized with 1 M Tris (pH 9) at a 1:5 ratio. Purified fractions were applied to protein concentrators (100 K MWCO; ThermoFisher) to change the buffer to PBS.

### ELISA

To quantify binding activity and specificity of the clones, scFv and IgG antibodies in lysates or purified fractions were analyzed by ELISA according to standard protocols. Briefly, Costar 96-well ELISA plates (Corning) were coated overnight at 4 °C with 50 µL of 1 µg/mL polysia-NCAM in PBS. After blocking in PBST (1% (v/v) Tween-20 in PBS) with 3% (w/v) milk for 2 h at room temperature, the plates were washed three times with PBST followed by incubation at room temperature for 1 h with serially diluted cell lysates or purified proteins. After washing three times with PBST, 100 µL of 1:1000-diluted anti-His (HRP) (Abcam) was added to each well and incubated for 1 h in the dark. The plates were then washed three times with PBST followed by the addition of 100 µL per well of 1-Step Ultra TMB (ThermoFisher). The reaction was quenched with 100 µL per well of 2 M H_2_SO_4_ and the Abs_450_ was measured in a multi-well plate reader (Tecan Spark).

### BLI analysis

The dissociation constants for selected scFvs and full-length IgGs were determined by BLI analysis. Briefly, 96-well plates containing samples diluted in kinetic buffer (1x PBS containing 0.2-μm filtered 0.1% (w/v) BSA and 0.02% (v/v) Tween 20) were analyzed using an Octet RH16 instrument (Sartorious) at 30 °C with shaking at 1,000 rpm. Kinetic analysis for scFv antibodies was performed using streptavidin biosensor tips (Sartorius 18–5020) as follows: (1) baseline: 30 s immersion in kinetic buffer; (2) loading: 500 s immersion in kinetic buffer supplemented with 0.2 μg/mL biotinylated polysia-NCAM; (3) baseline: 60 s immersion in kinetic buffer; (4) association: 600 s immersion in kinetic buffer solutions with varying scFv concentrations ranging from 35 to 500 nM; and (5) dissociation: 600 s immersion in kinetic buffer. Kinetic analysis of IgG antibodies was performed similarly except that biotinylated IgGs were immobilized on SA biosensor tips and immersed in kinetic buffer solutions with varying polysia-NCAM concentrations ranging from 2 to 134 nM. Kinetic data was analyzed using the Octet Analysis Studio software v12.2.2.26 (Sartorious).

### Cell binding assays

To assess binding specificity, we performed flow cytometric binding assays with human cancer cell lines, including SW2 and SH-SY5Y, that are highly polySia-positive ^63^. Briefly, cells were passaged at least three times before flow cytometry binding assays. On the day of the experiment, cells were trypsinzed and collected with media. The cells were centrifuged at 1,000 ×g and resuspended to 1×10^6^ cells/100 μL using PBSA (0.5% (w/v) BSA in PBS) and pipetted into round-bottom 96-well plates. Cells were washed three times with PBSA, then pelleted and resuspended in PBSA containing IgGs serially diluted from 15 μg/mL and incubated for 30 min at room temperature with constant agitation. Cells were washed three times with PBSA and resuspended in goat anti-human IgG-AF488 secondary antibody (ThermoFisher) at a 1:200 dilution for 30 min at room temperature in the dark with constant agitation. Cells were washed three times, resuspended in 200 μL of PBSA, and analyzed on a BD FACSMelody Cell Sorter (BD Biosciences). To remove polySia, SW2 cells were treated with 3 μg/mL endoN by adding the enzyme directly to culture media and incubating overnight.

### ADCC assay

Cryopreserved PBMCs (Cytologics, cat # 1105-C100) and SW2 cells were used to evaluate ADCC activity of all chAb735 constructs. PBMCs effector cells were thawed and rested in RPMI, supplemented with L-Glutamine and 10% FBS according to the manufacturer’s protocol. SW2 target cells were grown and maintained in high-glucose DMEM without phenol red, supplemented with 10% (v/v) FBS, penicillin (100 U/mL) and streptomycin (100 μg/mL). On the day of the experiment, SW2 cells were labeled with 25 μM calcein AM (Invitrogen) at a concentration of 10^5^ cells/mL culture medium at 37 °C and 5% CO_2_ for 1 h. Excess calcein was removed by washing with PBS and cells were resuspended in RPMI medium without phenol red, supplemented with 10% (v/v) FBS. Labeled SW2 cells were aliquoted in v-bottom 96-well plate (50 μL, 1×10^4^ cells/well). Ten-fold serial dilutions of polySia-specific chAbs were added to each well for the detection of experimental lysis. After antibody opsonization, 2×10^5^ PBMCs were added to each well at an effector-to-target cell ratio of 20:1 and incubated at 37 °C and 5% CO_2_ for 6 h. SW2 cells treated with wt chAb735 and chAb735^D158G/A230G^ in the absence of PBMCs or with an isotype chAb in the presence of PBMCs served as negative controls. The cell mixtures were centrifuged at 300 ×g for 5 min and the supernatant was transferred to flat-bottom 96-well plate. The released fluorescence was measured at 490/530 excitation/emission wavelengths using a multi-well plate reader (Tecan Spark). The specific ADCC was calculated using the following formula: ADCC (%) = (E–S)/(L–S)*100, where E is fluorescence with experimental antibody, and S is spontaneous fluorescence without antibody, and L is fluorescence obtained from the addition of a lysis buffer containing 1% (v/v) Triton-X 100.

### CDC assay

SW2 cells and baby rabbit complement were used to evaluate CDC activity of all chAb735 constructs. Briefly, cells were passed five times to ensure maximum population of viable cells before seeding in 96-well plates (Corning) at a cell density of 1×10^4^ cells/well. Cells were left incubating overnight at 37 °C and 5% CO_2_ for complete adhesion. Ten-fold serial dilutions of IgGs starting at 10 μg/ml were added in the presence of 10% (v/v) baby rabbit complement (MP Biomedicals) and incubated for 3 h. Cells treated with wt chAb735 and chAb735^D158G/A230G^ in the absence of complement or with an isotype chAb in the presence of complement served as negative controls. To measure cell viability, the CellTiter 96^®^ AQ_ueous_ One Solution Cell Proliferation Assay (Promega) was used according to manufacturer’s instructions. Briefly, 20 μL of CellTiter 96^®^ AQ_ueous_ One Solution Reagent was added to each well and further incubated for 3 h at 37°C and 5% CO_2_, after which Abs_490_ was measured. The specific cytotoxicity in each well was calculated using the following formula: cytotoxicity (%) = 100 x (B - E)/(B – L), where E is the absorbance with experimental antibody, and B is the absorbance without antibody but with the same concentration of serum, and L is the absorbance obtained from the cells lysed with lysis buffer.

### Statistical analysis

Statistical significance between groups was determined with unpaired two-tailed Student’s *t*-test using GraphPad Prism software for MacOS (version 9.4.1). Statistical parameters including the definitions of *n*, *p* values, and SDs are reported in the figures and corresponding figure legends.

### Data availability

All data generated or analyzed during this study are included in this article and its Supplementary Information/Source Data file that are provided with this paper.

## Acknowledgments.

We thank Dr. Dane Wittrup (MIT) for providing plasmid pCT-CON. Dr. Gaurang Bhide and Dr. Karen Colley (University of Illinois at Chicago) for the SW2 cell line as well as purified endoN used in this work. This work was supported by the Defense Threat Reduction Agency (grants HDTRA1-15-10052 and HDTRA1-20-10004 to M.P.D.), the National Science Foundation (grants CBET-1605242, CBET-1936823, and MCB-1413563 to M.P.D.), the National Institutes of Health (grants R01 GM137314- 01 to M.P.D.), and a Cornell Fleming Graduate Scholarship (to N.L.-B.).

## Author Contributions

W.W. designed research, performed research, analyzed data, and wrote the paper. M.B. and N.L.-B. designed research, performed research, and analyzed data. M.P.D. designed and directed research, analyzed data, and wrote the paper. All authors read and approved the final manuscript.

