## Supplementary Figures 1-8 for "Engineering affinity-matured variants of an anti-polysialic acid monoclonal antibody with superior cytotoxicity-mediating potency"

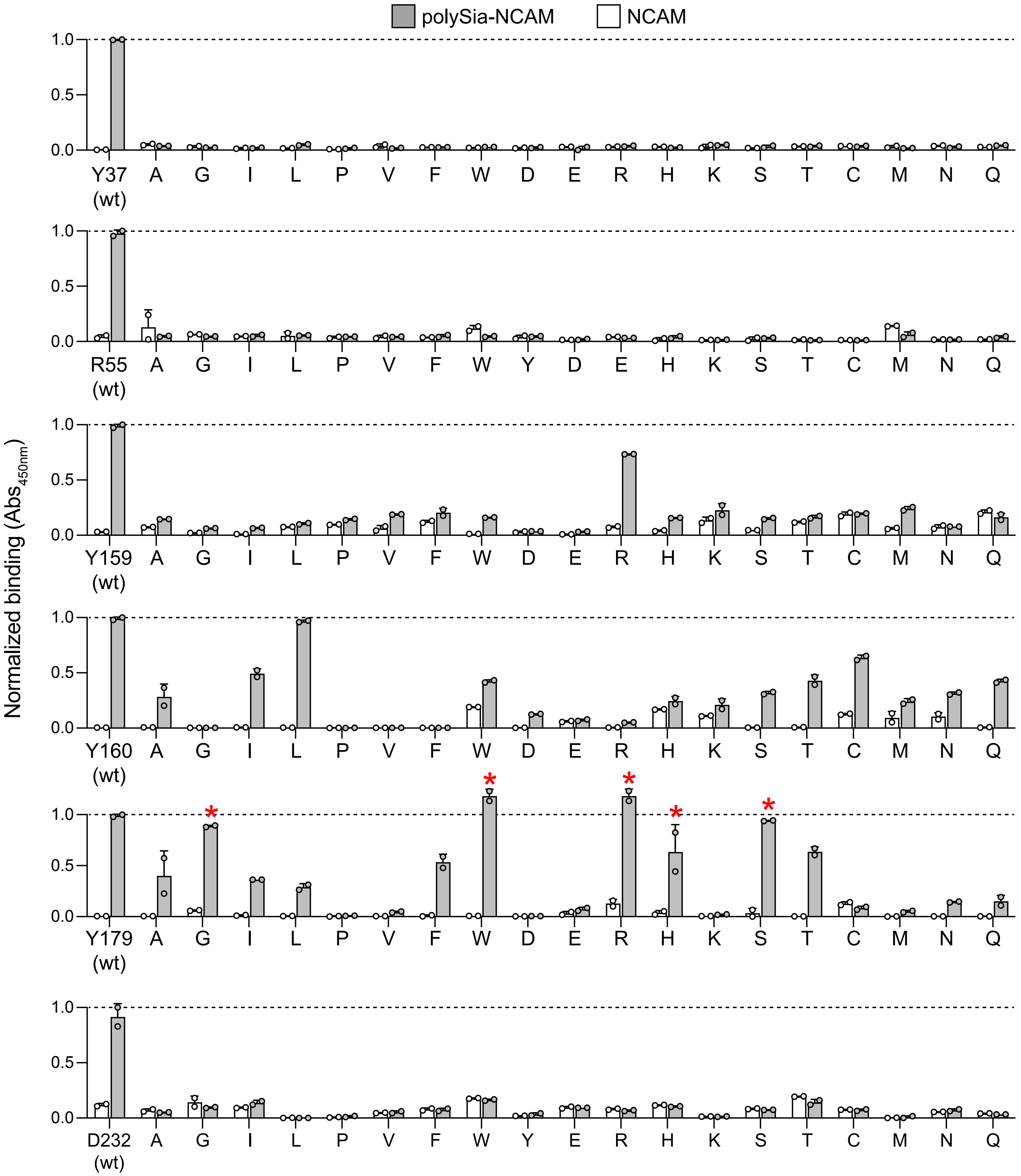


**Supplementary Figure 1. Relative importance of residues that make direct contact with polySia antigen.** Binding analysis of six direct contact residues (L:Tyr-37, L:Arg-55, H:Tyr-159, H:Tyr-160, H:Tyr-179, and H:Asp-232) in scFv735. Each contact residue was subjected to SSM, resulting in a set of 114 site-directed variants of scFv735 in which all 19 amino acid substitutions were introduced at each of the six positions. All 114 scFv735 variants and wt scFv735 were expressed from plasmid pMAB in *E. coli* SHuffle T7 Express cells and collected as cell lysates. Binding activity in cell-free lysates was quantified by ELISA using either polySia-NCAM (gray bars) or endoN-treated NCAM (white bars) as immobilized antigen. An equivalent amount of total protein was loaded in each well. Data are average of biological replicates ± SD and are normalized to signals obtained for wt scFv735. Red asterisks denote variants selected for further analysis.

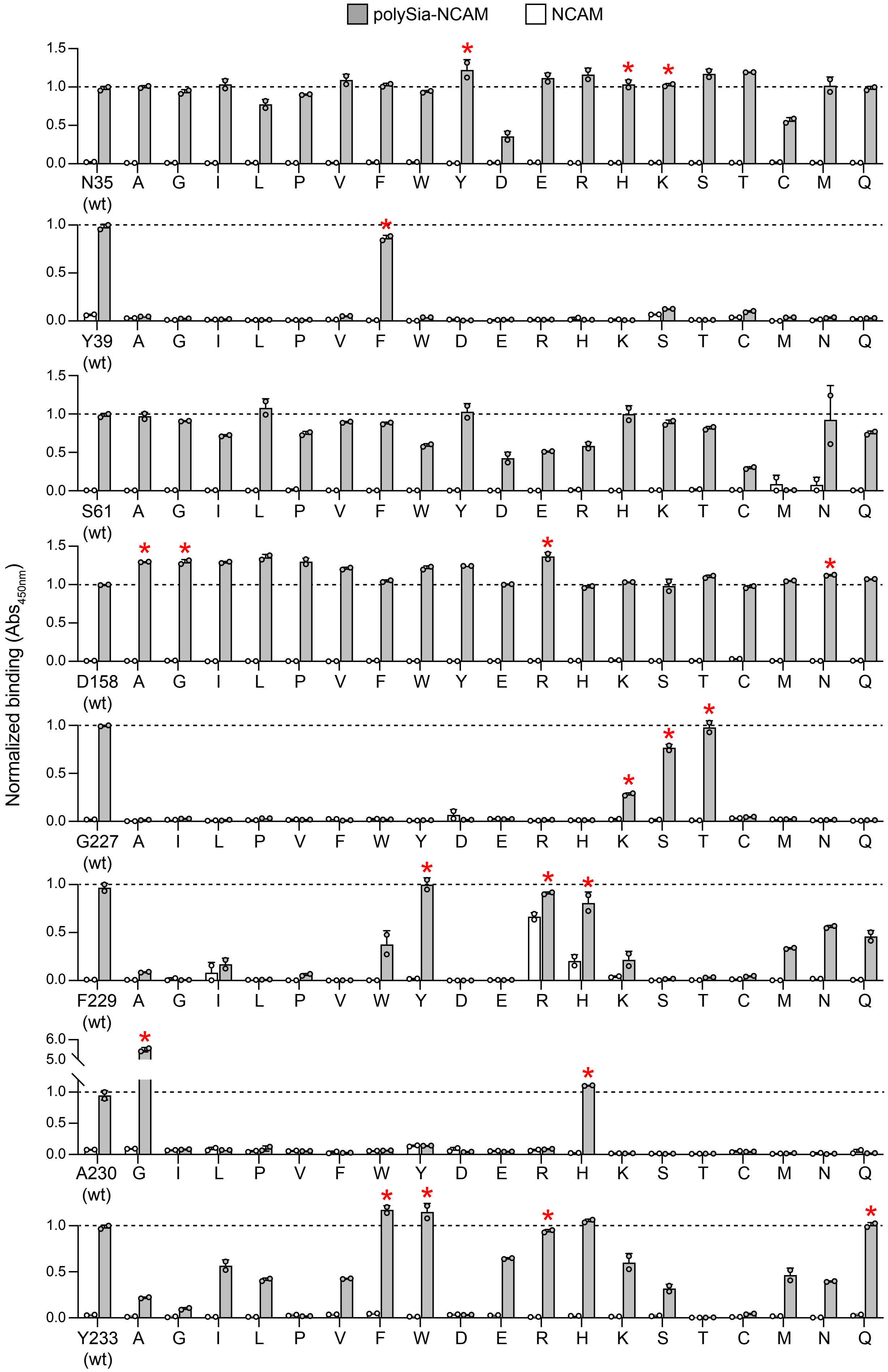


**Supplementary Figure 2. Relative importance of residues that make direct contact with polySia antigen.** Binding analysis of eight indirect contact residues (L:Asn-35, L:Tyr-39, L:Ser-61, H:Asp-158, H:Gly-227, H:Phe-229, H:Ala-230, and H:Tyr-233) in scFv735. Each contact residue was subjected to SSM, resulting in a set of 152 site-directed variants of scFv735 in which all 19 amino acid substitutions were introduced at each of the six positions. All 152 scFv735 variants and wt scFv735 were expressed from plasmid pMAB in *E. coli* SHuffle T7 Express cells and collected as cell lysates. Binding activity in cell-free lysates was quantified by ELISA using either polySia-NCAM (gray bars) or endoN-treated NCAM (white bars) as immobilized antigen. An equivalent amount of total protein was loaded in each well. Data are average of biological replicates ± SD and are normalized to signals obtained for wt scFv735. Red asterisks denote variants selected for further analysis.

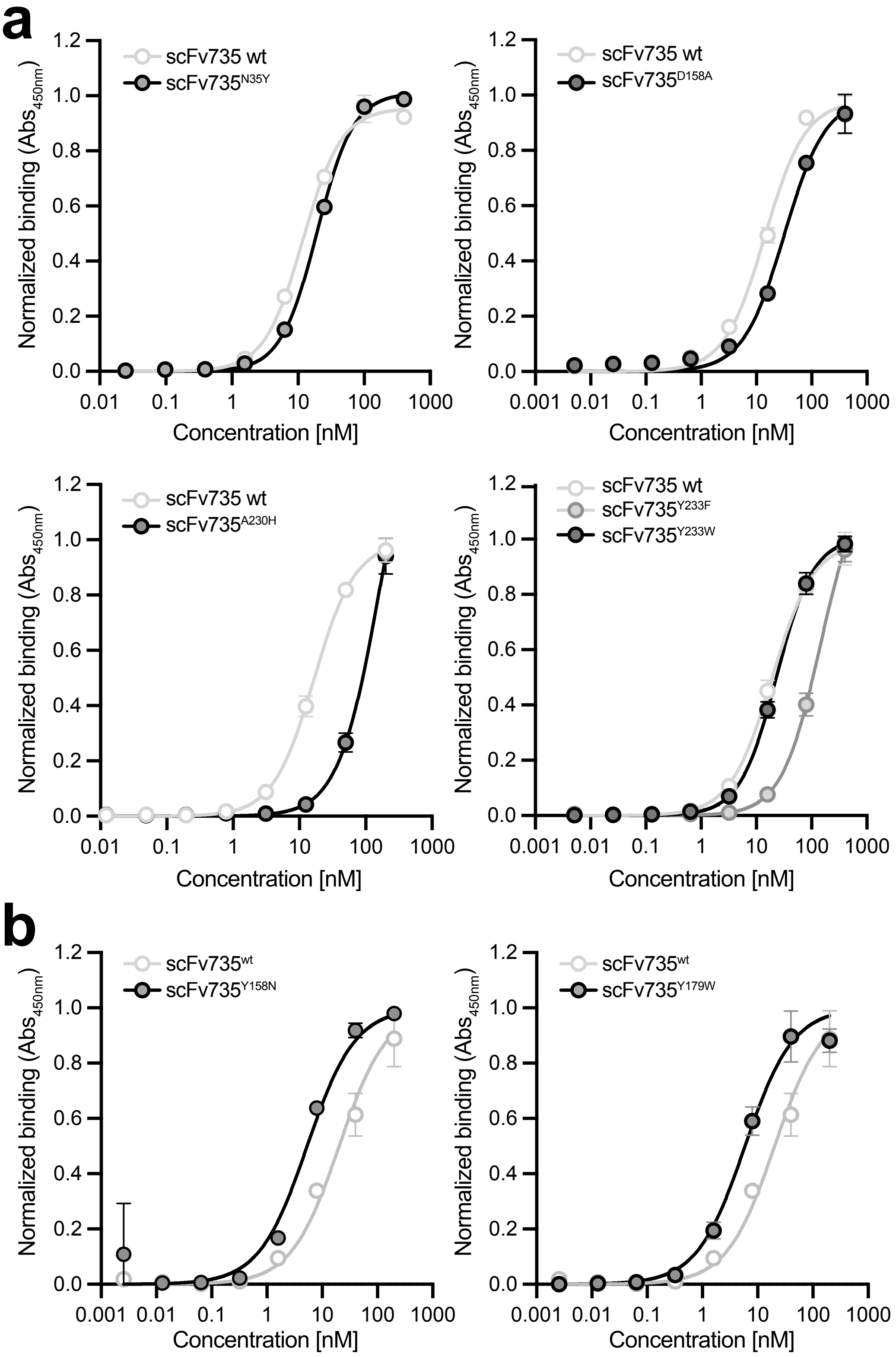


**Supplementary Figure 3. Binding curves for representative single-site variants from SSM screening.** ELISA analysis of select site-directed mutants exhibiting (a) non-improved or (b) improved binding activity (gray circles) relative to wt scFv735 (white circles). Each variant was expressed from plasmid pMAB in *E. coli* strain SHuffle T7 Express cells and purified from cell-free lysate by Ni-NTA affinity chromatography. PolySia-NCAM was used as immobilized antigen. Data are average of biological replicates ($n = 3$) ± SD.

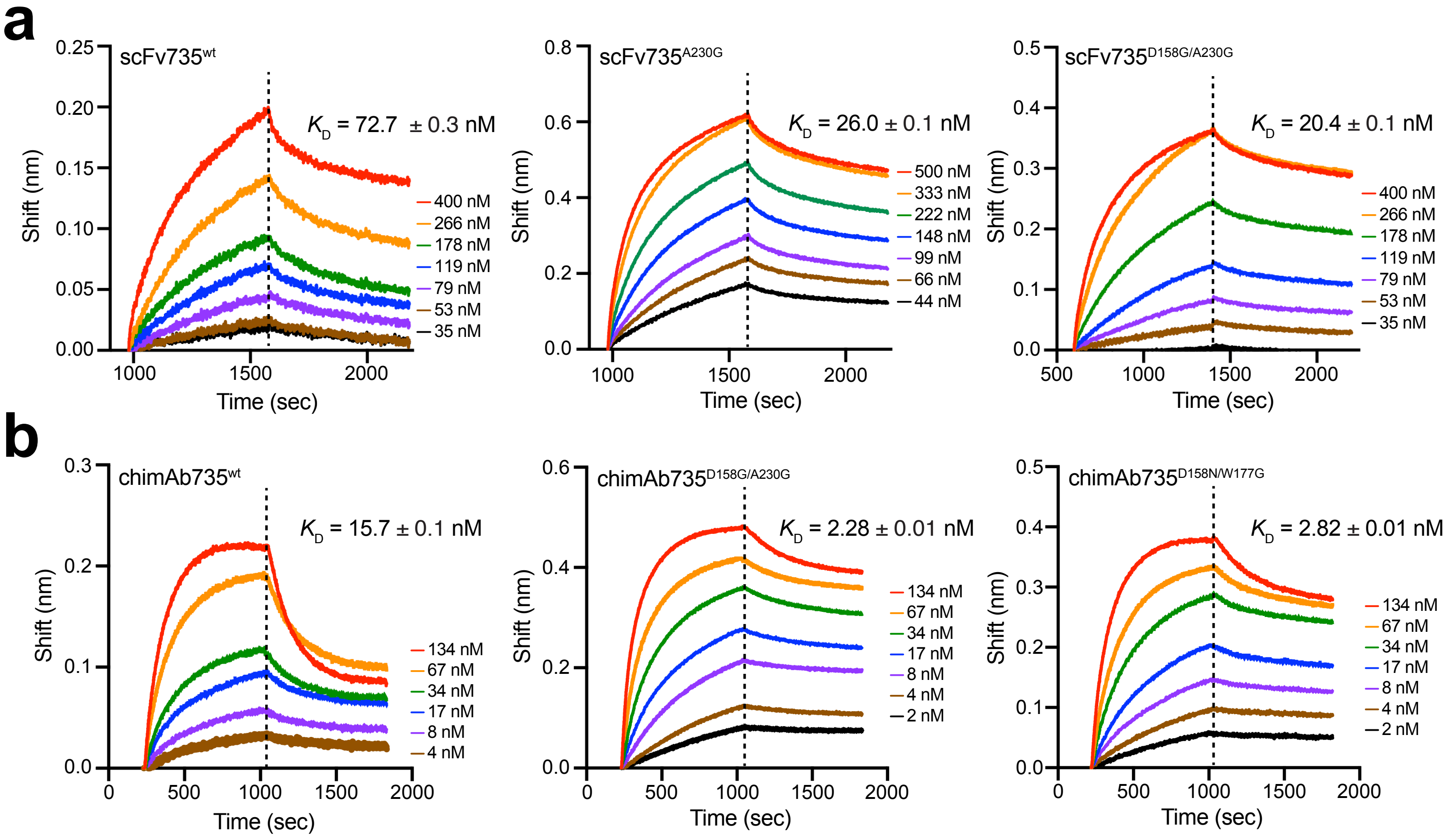


**Supplementary Figure 4. Binding affinity for representative improved scFv735 and chAb735 variants.** BLI analysis to quantify kinetic binding constants ($k_{on}$, $k_{off}$) and equilibrium binding constant ($K_D$) for the interaction between polySia-NCAM and purified (a) scFv735 and (b) chAb735 antibodies. Biotinylated polySia-NCAM was immobilized on streptavidin (SA)-coated sensors and subsequently used to bind scFvs or chAbs in solution. Response data are representative of replicate BLI experiments. Binding constants were determined by fitting response data to Langmuir model in Octet Analysis Studio (see Table 1).

**Mutations in NNK-derived variants (all in CDR H2):**

H2Lib1-3: **W177R I178V Y179F G181K**

H2Lib1-4: **W177R I178R P180G G181K S182G**

H2Lib2-2: **W177S G181R S182T G183S N184R T185G K186R**

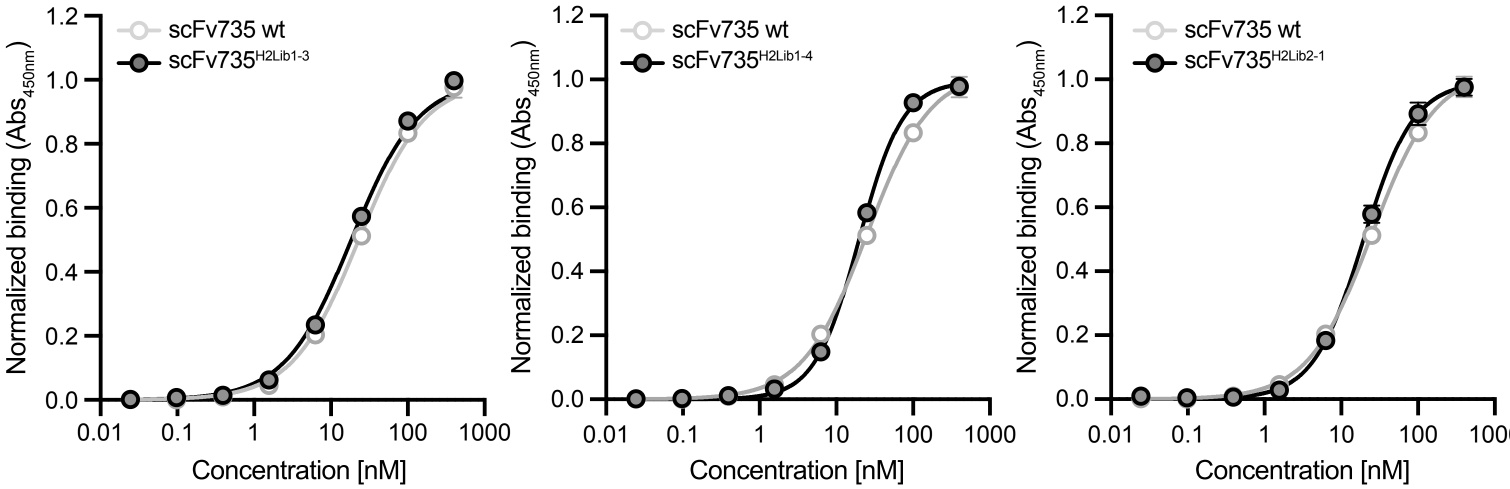


**Supplementary Figure 5. Binding curves for improved NNK variants.** ELISA analysis of select NNK mutants of scFv735 expressed from plasmid pMAB in *E. coli* strain SHuffle T7 Express cells and purified from cell-free lysate by Ni-NTA affinity chromatography. Representative ELISA binding curves comparing wt scFv735 (white circles) with improved variants (gray circles). PolySia-NCAM was used as immobilized antigen. An equivalent amount of total protein was loaded in each well. Data are average of biological replicates ($n$ = 3) ± SD.

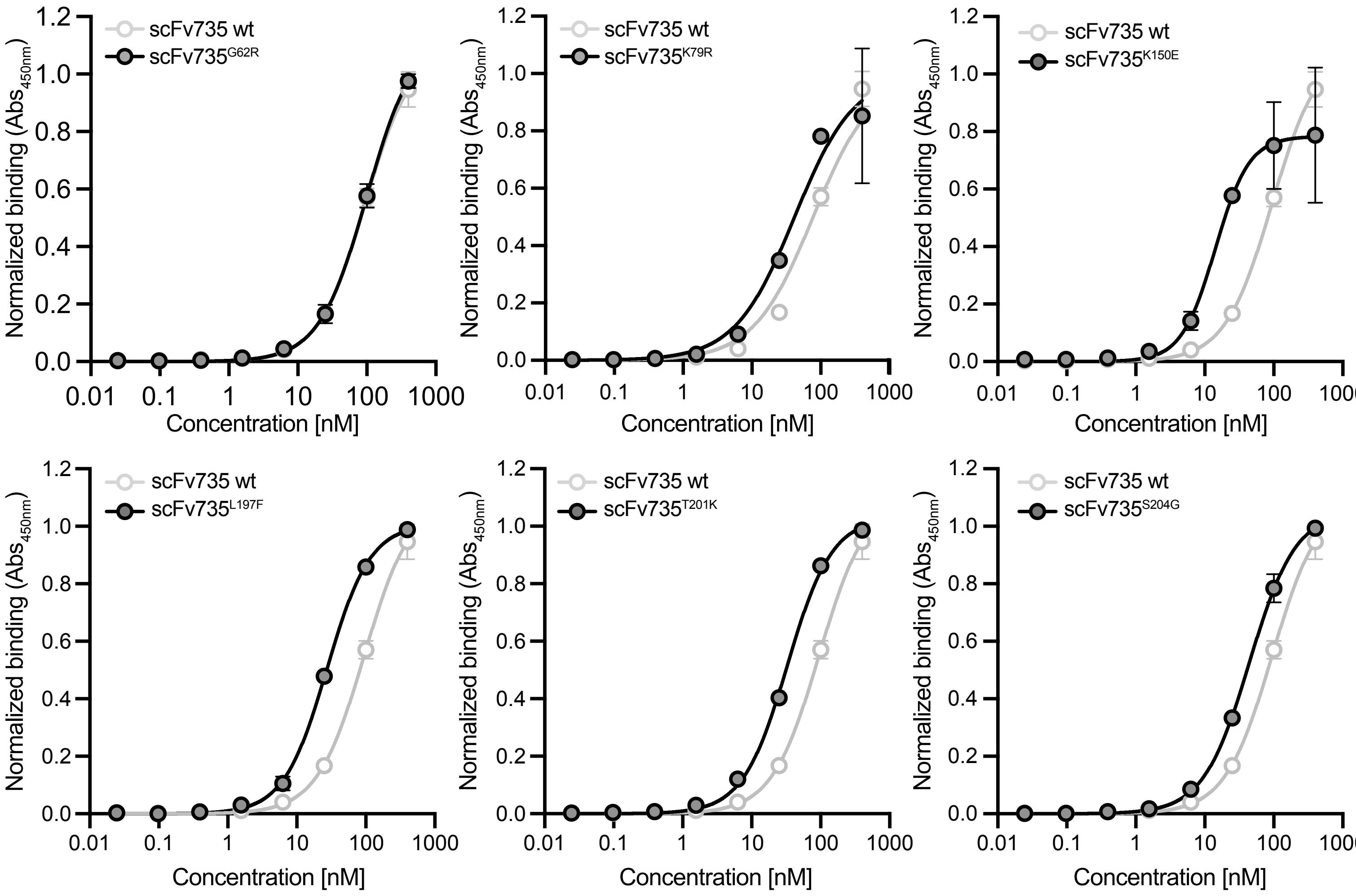


**Supplementary Figure 6. Binding curves for improved epPCR variants.** ELISA analysis of select epPCR mutants of scFv735 expressed from plasmid pMAB in *E. coli* strain SHuffle T7 Express cells and purified from cell-free lysate by Ni-NTA affinity chromatography. Representative ELISA binding curves comparing wt scFv735 (white circles) with improved variants (gray circles). PolySia-NCAM was used as immobilized antigen. An equivalent amount of total protein was loaded in each well. Data are average of biological replicates ($n$ = 3) ± SD.

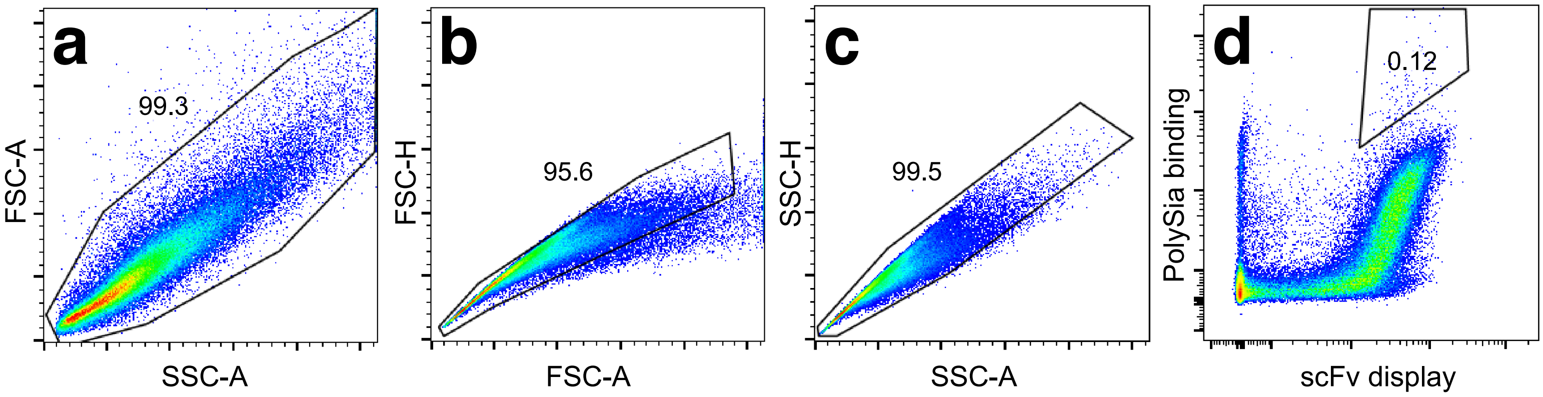


**Supplementary Figure 7. Gating strategy for FACS-based isolation of polySia-specific scFv clones.** (a-d) Representative flow cytometry scatter plots showing the gating strategy used to isolate polySia-binding clones from yeast surface display libraries. A total of at least 10,000 events were captured on a BD FACSMelody instrument and data were analyzed using FlowJo software. Typical forward scatter versus side scatter plots are depicted in (a), showing the first gating step to eliminate cell debris. Next, sorting of single cells was performed by gating based on (b) forward scatter (FSC) signals and (c) side scatter (SSC) signals. Finally, in (d), about 0.1% of yeast cells were sorted based on high signals correpsonding to scFv expression/display on the cell surface and polySia binding. These signals were generated by labeling with rabbit anti-Myc tag antibody followed by Alexa Fluor 647-conjugated goat anti-rabbit IgG antibody for scFv display and with biotinylated polySia-NCAM followed by streptavidin-Alexa Fluor 488 conjugate for polySia binding.

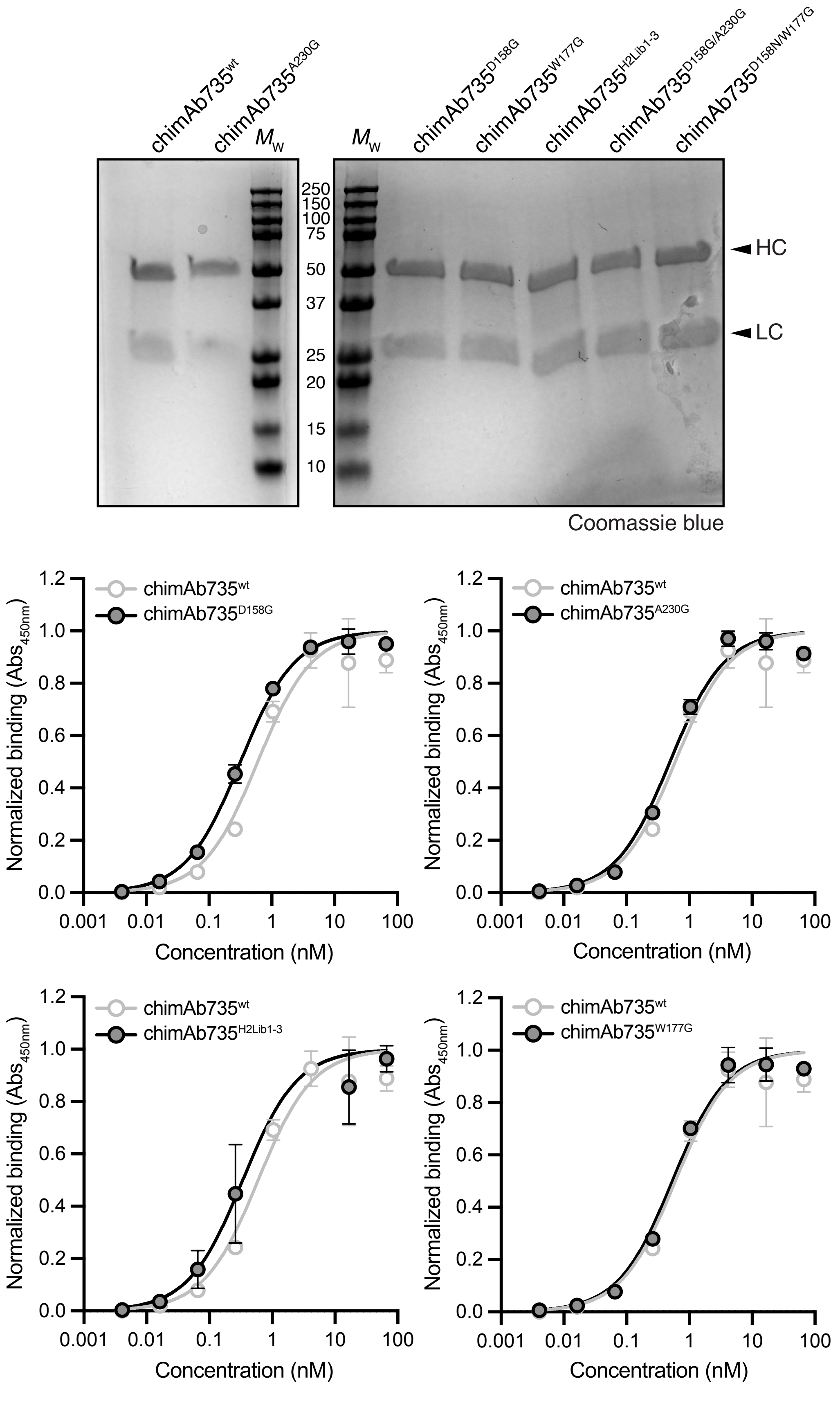


**Supplementary Figure 8. Binding curves for representative improved chAb735 variants.** ELISA analysis of select chAb735 mutants expressed from plasmid pVITRO1 in Freestyle 293-F cells and purified from culture supernatants by protein A affinity chromatography. Representative ELISA binding curves comparing wt chAb735 (white circles) with improved variants (gray circles). PolySia-NCAM was used as immobilized antigen. An equivalent amount of total protein was loaded in each well. Data are average of biological replicates ($n = 3$) ± SD.
